# Pregnancy and postpartum dynamics revealed by millions of lab tests

**DOI:** 10.1101/2023.05.11.540359

**Authors:** Alon Bar, Ron Moran, Netta Mendelsohn-Cohen, Yael Korem Kohanim, Avi Mayo, Yoel Toledano, Uri Alon

**Affiliations:** Dept. Molecular Cell Biology, Weizmann Institute of Science; Rehovot, 7610001, Israel; Dept. of Computer Science and Applied Mathematics, Weizmann Institute of Science; Rehovot, 7610001, Israel; Department of Immunobiology, Yale University School of Medicine, New Haven, CT; Division of Maternal Fetal Medicine, Helen Schneider Women’s Hospital, Rabin Medical Center; Petah Tikva, 4941492, Israel

## Abstract

Pregnancy and delivery involve dynamic alterations in many physiological systems. However, the physiological dynamics during pregnancy and after delivery have not been systematically analyzed at high temporal resolution in a large human population. Here we present the dynamics of 76 lab tests based on a cross-sectional analysis of roughly 41 million measurements from over 300,000 pregnancies. We analyzed each test at weekly intervals from 20 weeks preconception to 80 weeks postpartum, providing detailed temporal profiles. About half of the tests take three months to a year to return to baseline during postpartum, highlighting the physiological load of childbirth. The precision of the data revealed effects of preconception supplements, overshoots after delivery and intricate temporal responses to changes in blood volume and renal filtration rate. Pregnancy complications – gestational diabetes, pre-eclampsia and postpartum hemorrhage – showed distinct dynamical changes. These results provide a comprehensive dynamic portrait of the systems physiology of pregnancy.

## Introduction

During pregnancy, the mother undergoes physiological changes that support fetal growth and development. The cardiovascular, respiratory, renal, gastrointestinal, skeletal, metabolic, endocrine and immune systems are all affected by fetal demand and massive endocrine secretion by the placenta(*1*–*4*). Increased demand for oxygen and nutrients causes an increase in cardiac output and up to 50% growth in blood volume(*1*). The kidneys increase the glomerular filtration rate by 50%, leading to increased urine production(*1*). The immune system is modulated to prevent rejection of the fetus, and coagulation and red blood cells show marked changes (*1, 2, 5*). Metabolism shifts to increased insulin resistance and lipid production to supply energy for fetal growth (*6*).

Delivery marks a profound change, as the fetus and placenta exit the body and abruptly cease their metabolic and endocrine effects. The mother undergoes a series of adaptations in which various physiological systems recover with different timescales - from hours to months(*7*). Pregnancy and postpartum periods have an increased risk of complications including gestational diabetes, postpartum hemorrhage, anemia, depression and eclampsia(*8*).

Understanding healthy physiology and pathology is essential for both advancing basic science and as a baseline for treatment. This understanding of the physiological changes during pregnancy and postpartum requires precise temporal data on numerous physiological parameters. However, existing studies have a limited number of participants, consider only a few parameters, and have low temporal resolution, typically of one time point per trimester (*9, 10*). Knowledge is even more sparse in the postpartum period in which a single time point is usually measured. Meta-analyses have collected these smaller studies to construct normal ranges for tests in each trimester (*9, 10*). Altogether, our knowledge of the physiological time course is thus limited to low temporal resolution. Here we harness a large national health record database (*11*) to study over 300K pregnancies in terms of 76 major lab tests, totaling over 40 million tests. These test results are cross-sectionally analyzed at weekly time intervals, and we present this information as a resource. We identify global dynamical trends in healthy pregnancies and in pregnancy complications.

## Results

### Lab tests throughout pregnancy and postpartum

We obtained data from Clalit Healthcare, the largest health maintenance organization (HMO) in Israel, with over 5 million members as of 2024, with broad socioeconomic and ethnic demographics (*11, 12*). The HMO recorded about half of the pregnancies in Israel between 2003-2020. We analyzed 313,501 pregnancies of females aged 20 to 35. The fraction of first pregnancies in the dataset is 47% **(Methods)**. The mean and median gestational week at delivery is 39 weeks. Preterm deliveries (<37 weeks) are 12% **(Methods)**. C-sections account for 16.4% of deliveries. This is an all-comers dataset, with stillbirths and multiple deliveries excluded. Demographic characteristics of the participants are presented in **Table 1**. We removed test values from individuals with disease codes or medications that statistically affect each test (**Methods**).Thus, we consider the dataset to include only healthy pregnancies.

**Table 1.** Age and BMI of the study population. Due to privacy concerns, other demographics were not available. Age statistics are for all participants who had any of the 76 analyzed tests in the indicated week. Due to the cross-sectional nature of the dataset each test and weekly interval is drawn from a different sub cohort of the study population. (IQR: interquartile range).

|  | 60 weeks before delivery | At delivery | 80 weeks postpartum |
| --- | --- | --- | --- |
| **Age (mean, IQR)** | 28.1 (25.1-31.3) | 28.2 (25.3-31.4) | 29.3 (26.6-32.2) |
| **BMI (mean, IQR)** | 24.4 (20.7-27.0) | 28.8 (25.1-31.9) | 26.0 (21.0-29.4) |

In the period of 60 weeks before delivery to 80 weeks after delivery, we identified 41,324,910 test values from 110 lab tests **(table S1, fig. S1)**. We filtered out 35 tests due to high noise and/or low number of measurements **(Methods) and** retained for analysis 76 tests with a total of 40,465,393 measurements. Each test had between 34,808 and 1,601,625 total measurement values. Our ethical agreement precluded longitudinal analysis of individual pregnancy trajectories. We therefore performed cross-sectional analysis - we aggregated each test over weekly intervals and analyzed them for summary statistics **(Methods)** including mean, median and the (5, 10, 25, 50, 75, 90, 95) percentiles. For clarity we highlight one test, alkaline phosphatase, in **Fig. 1B** and show all 76 profiles in **Fig. 1C**. Tests arranged by physiological systems are discussed in detail in the supplementary text and **fig. S2**.

**Fig. 1.**
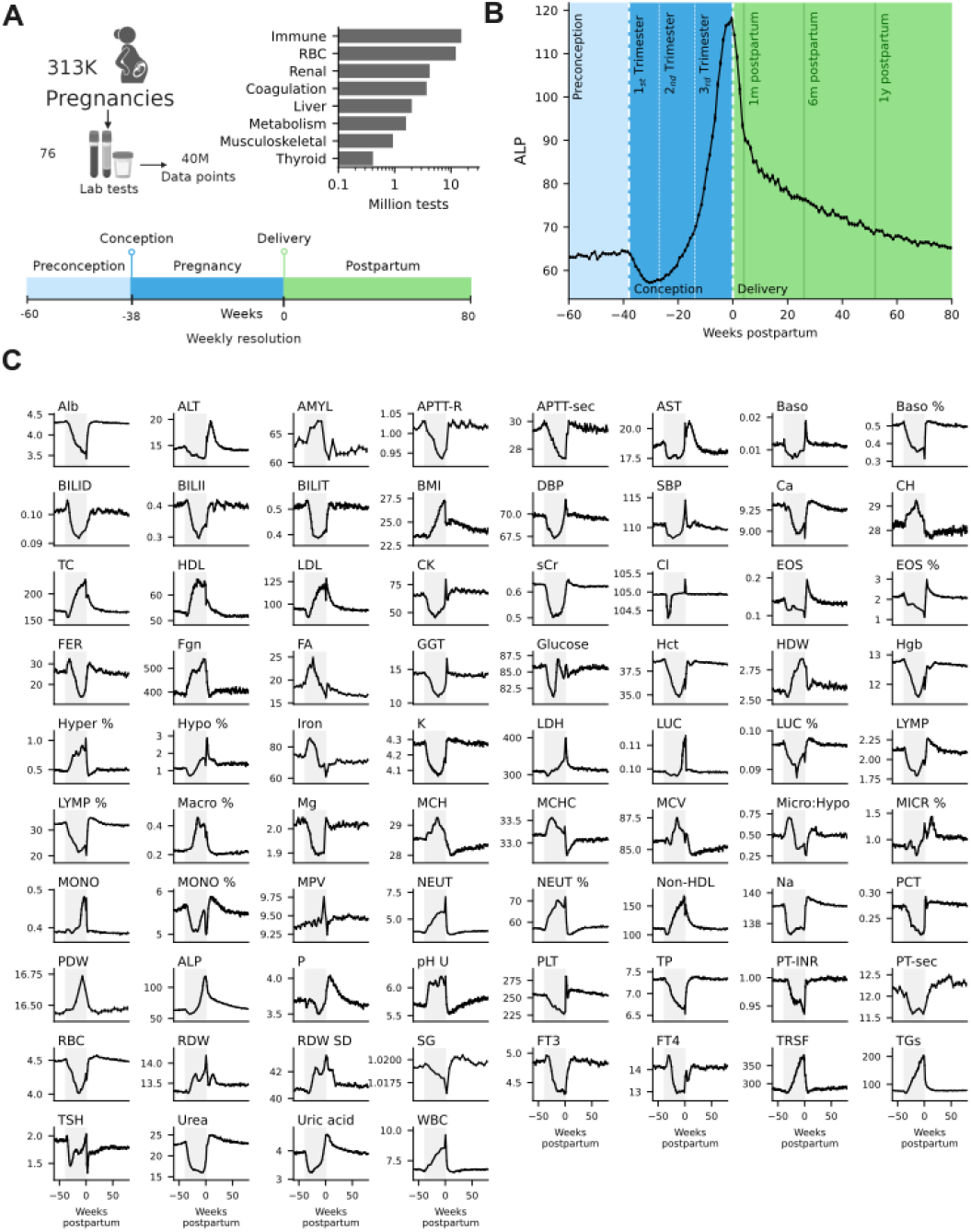
Dataset of lab tests over preconception, pregnancy and postpartum. A) Schematic overview of the dataset. B) Alkaline phosphatase test (mean of the quantile-transformed values, see **Methods**) over 140 weeks, with marks for 22 weeks before conception, 38 weeks of pregnancy and 80 weeks after delivery. Error bars are standard error of the mean. Units are the standard for the test (IU/L). C) 76 test values over 140 weeks, the period of pregnancy is in gray. For units see table S1. Graphs in some panels are smoothed for visualization **(Methods)**.

### Dynamic variation in pregnancy scales with homeostatic variation

All 76 mean test values varied across pregnancy and postpartum with a dynamic range of a few percent to hundreds of percent for different tests. The dynamic range of each test rises with its variation within the reference population - tests that vary widely between individuals also show large dynamic ranges during pregnancy and postpartum (**Fig. 2A**, Pearson correlation coefficient=0.72, *p* < 10^−5^) **(Methods)** with a linear relation. Thus, homeostatic processes seem to remain mostly within their physiological range. In the following analysis we therefore use quantiles scores relative to a non-pregnant population. Quantile scores varied by 5 to 52 percentile points relative to reference, with a mean of 29 percentiles **(Fig. 2B)**.

**Fig. 2.**
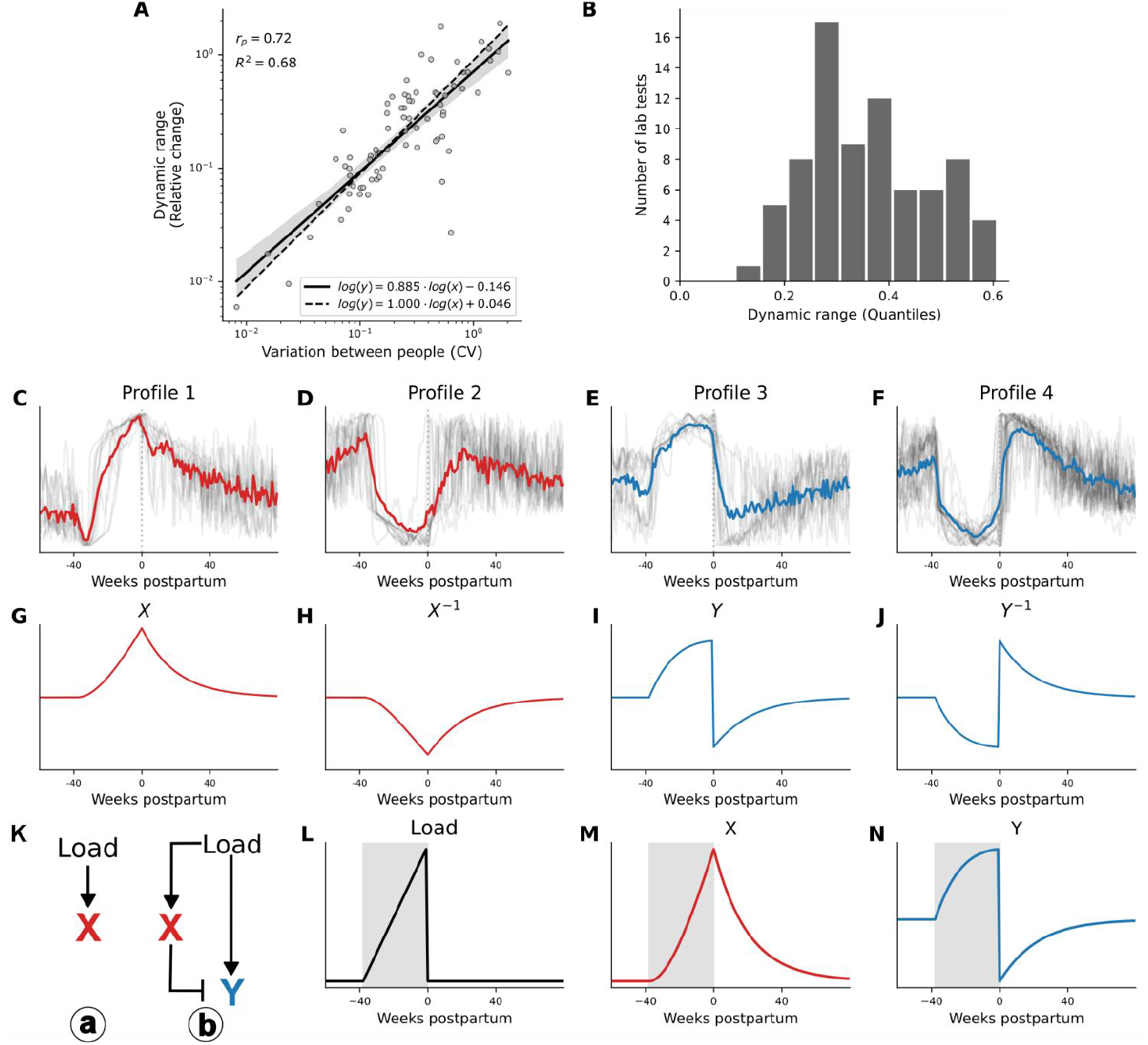
Lab test dynamics vary during pregnancy and postpartum and can show overshoots and undershoots. **A)** Relative dynamic range across the study period of back-transformed mean test values (Methods), log scale with base 10 on both axes. Dynamic range (max-min)/(preconception average) rises with the coefficient of Variation (CV) of the test value in the reference population. r_p_ is the Pearson correlation. The dashed line within the 95% confidence interval is the linear model, R^2^=0.61. **B)** Histogram of dynamic range of quantile scores of the 76 tests. **C-F)** Four clusters of ranked test dynamical profiles.

### Dynamics show overshoots and undershoots after delivery

To understand the observed dynamics, we clustered tests according to their temporal profiles **(Methods, fig. S4) (Fig. 2C-F)**. We found four clusters, which define four profiles. Two of the profiles change direction at delivery: Profile 1 rises during pregnancy and drops postpartum, and profile 2 is its mirror image, declining in pregnancy and rising postpartum. Profiles 3 and 4 show overshoots or undershoots at delivery followed by a return to steady state close to the preconception levels. To understand the origin of the overshoots we consider canonical physiological mechanisms **(Methods)**. Pregnancy exerts a load on physiological variables to meet the needs of the mother and developing fetus. This load pushes physiological variables away from their normal set points or adjusts new set points that reflect physiological priorities (*13, 14*). Upon delivery this load is suddenly relaxed **(Fig. 2L)**.

Variables without overshoot can be explained by first-order recovery to baseline **(Fig. 2K(a))**. Variables are pushed by the load away from steady state during pregnancy and recover postpartum with a characteristic timescale which follows the relaxation of the gestational load **(Fig. 2K(a),M)**. In this scenario, no overshoot occurs.

In contrast, an overshoot can occur when there exists an additional, slowly varying compensation mechanism **(Fig. 2K.2,N)**, modeled here by an incoherent feed forward loop circuit (*15, 16*). During pregnancy the compensatory mechanism intensifies to keep the variable from moving too far from its setpoint. Upon delivery the load is suddenly reduced but the compensation mechanism is still strong - causing overcompensation that induces an overshoot. Return to baseline of the variable is governed by the return of the compensation mechanism. An example is the slow growth of the thyroid gland during pregnancy that compensates for changes in the demand for thyroid hormones (*17*).

Other models can also provide the observed temporal shapes. For example, a load that grows during pregnancy and reduces during postpartum can provide alternative explanations for profiles 1 and 2. Delivery itself can serve as another source for rebound dynamics by forcing a sharp pulse in the opposite direction. However, such models require more complex control mechanisms with more parameters.

In gray are ranked individual tests; colored lines are their mean. **G-H)** Theoretical profiles for first-order response I-J) Theoretical profiles for a system with a slow compensatory mechanism. **K) (a)** Circuit in which the load of pregnancy affects test X as a first-order system. **(b)** Circuit in which load affects test Y with X as a compensatory system. Mathematical analysis can be found in the supplementary text **L)** Pregnancy load in the theoretical model rises in pregnancy and drops abruptly at delivery **M)** First order system X responds with no undershoot **N)** Compensated system Y shows an undershoot and a rebound effect.

### Physiological changes show slow postpartum recovery

To study the global temporal trajectories, we reduced dimensionality using principal component analysis on the 76 tests at all 140 week-intervals. The first two principal components capture 88% of the variation. The trajectory shows hysteresis - tests change during pregnancy and return to baseline via a different trajectory postpartum **(Fig. 3A)**.

**Fig. 3.**
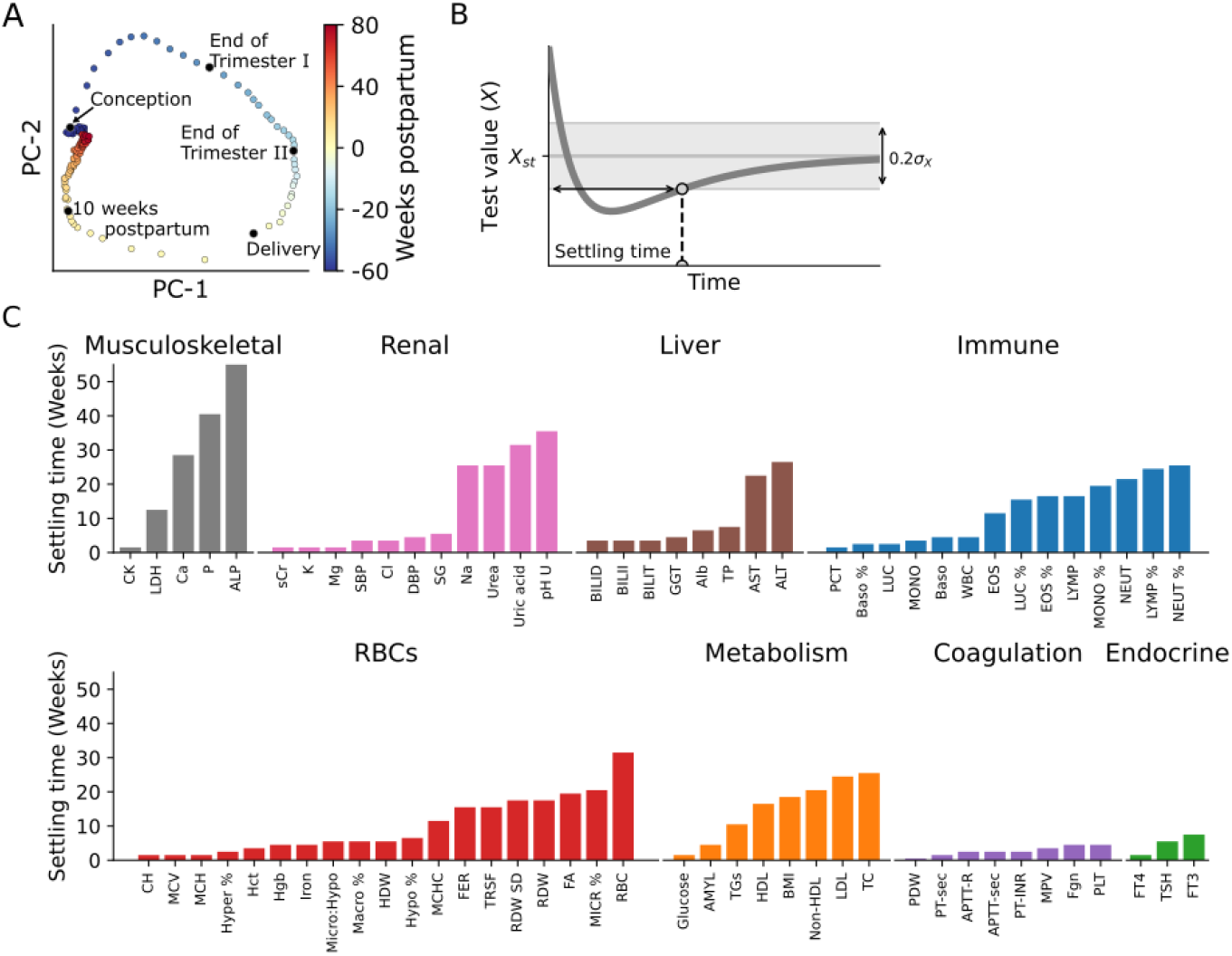
Postpartum recovery times of tests range between days and a year. **A)** Dimensionality reduction of test mean values as a function of time using PCA shows that the trajectory during pregnancy (blues) differs from the postpartum trajectory (orange and red). Gestational week progression is in the clockwise direction. **B)** Settling time is defined as the time after which the test remains within 0.2 standard deviations of its baseline. **C)** Settling time of the 76 tests in the dataset.

Postpartum adaptation has two main phases which are apparent on PC1 **(Fig. 3A)**. Most changes take place in the 10 weeks after delivery, followed by a prolonged return to steady state. Many tests take months to return to baseline after delivery. To quantify the time to return to baseline we use a measure from control theory called “settling time”. The settling time is defined by the time after which the test remains within a small margin (here 0.2 standard deviations) of its postpartum baseline **(Fig. 3B, Methods)**.

Approximately 42% (32/76) of the tests have long settling times that exceed 10 weeks **(Fig. 3C)**. Among these are liver functions aspartate transaminase (AST) and alanine transaminase (ALT) that take about half a year to recover, metabolic factors such as cholesterol, and alkaline phosphatase (ALP) that settles only after about a year **(Fig. 3C)**. Approximately 46% (35/76) of the tests settle rapidly within the first month. This includes all coagulation tests **(Fig. 3C)**. The remaining ∼12% of the tests (9/76) settle between one month to 10 weeks after delivery **(Fig. 3C)**.

Slow return to baseline can arise from several factors. Metabolism is affected by BMI that settles over months(*18*). Breastfeeding may also affect some tests, such as ALP, calcium, phosphate, PTH and prolactin(*19*). The dataset does not include information on who breastfed. 90% of Israeli neonates are breastfed for a mean duration of about 70 days. The rate of exclusive breastfeeding drops to about 60% at 2 months and to 20% at 6 months after birth (*20*).

Notably, several tests do not return to their preconception baseline, including elevated levels of the inflammation marker CRP, reduced thyroid stimulating hormone TSH and reduced mean cell hemoglobin MCH and iron.

### Health style behaviors are reflected in preconception dynamics

We noticed that about a third of the tests (24/76) show significant dynamical trends *before* conception, in the period of 60 to 38 weeks before delivery **(Fig. 4ABC) (Methods)**. One of the strongest changes is a rise in folic acid **(Fig. 4A)**. Folic acid supplements are taken in the months before conception by about half of the relevant population (*21, 22*).

**Fig. 4.**
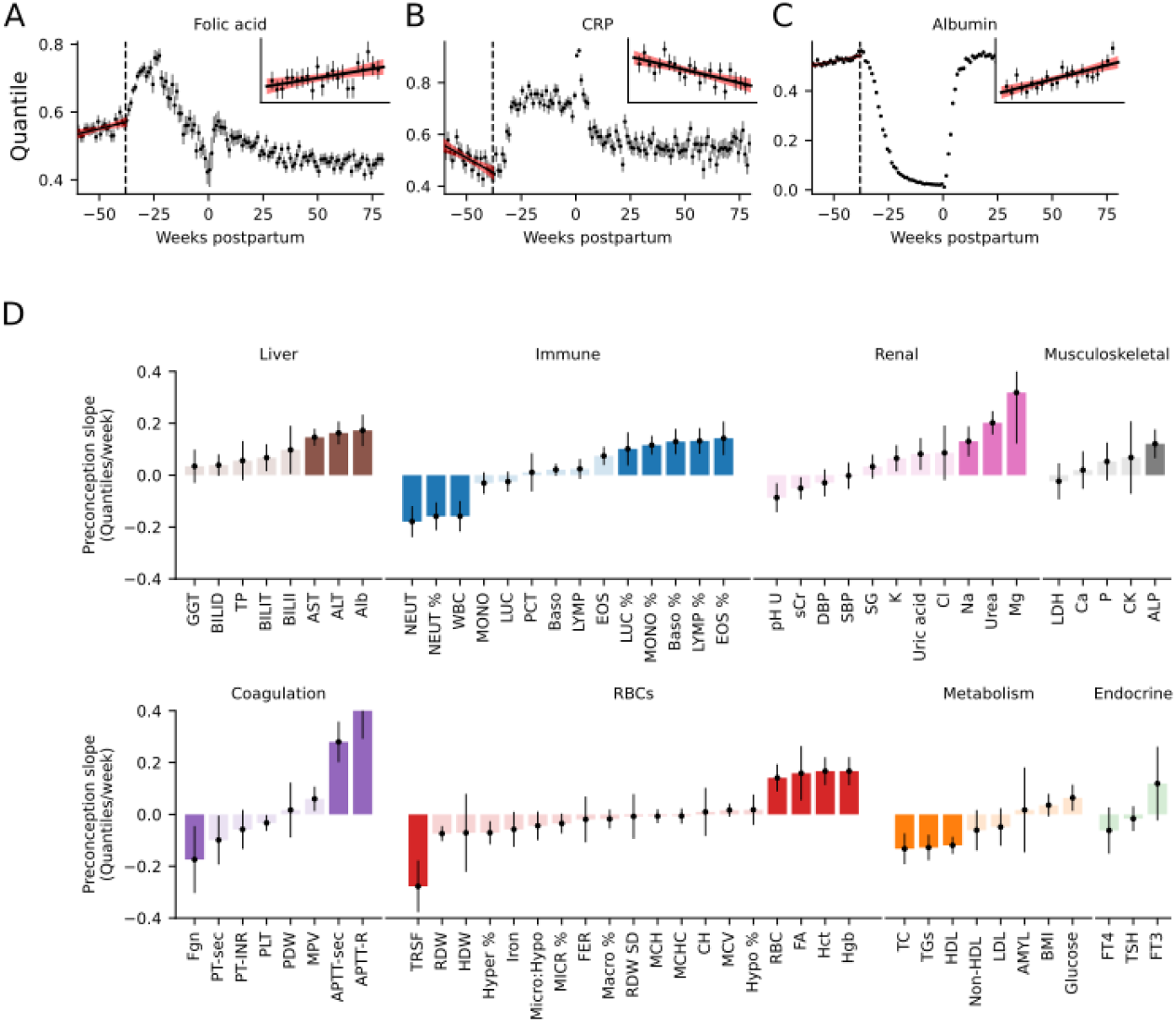
Tests affected by health behaviors show preconception dynamics. **A-C)** Folic acid, CRP and albumin are examples of a strong preconception change. Mean conception time is indicated by a dashed line. The inset highlights the preconception period, red is regression with 95% confidence intervals. **D)** Temporal slope (quantile change per week) of test values during preconception from linear regression. Tests are arranged by physiological system, tests in bold color have significant nonzero slope (>0.1 in absolute value and p<0.05 adjusted for multiple comparisons, Benjamini-Hochberg). Error bars are 95% confidence intervals of slope.

Supplements such as folic acid and vitamin B-12 can exert physiological effects on other test values. These changes include a reduction in CRP **(Fig. 4C)**, an increase in albumin **(Fig. 4D)**, positive effects on anemia, anticoagulative effects and lowering of lipids (*23*–*27*).

Some of the changes seen in preconception are not easily attributed to known effects of supplements. This includes changes in immune cell counts, ALT, AST, Na, Urea and Urine pH. One possibility is that these tests are affected by yet unknown mechanisms by supplements, or that they are affected by other preconception health behaviors such as reduced rates of smoking, alcohol consumption and improved diet (*28*).

We conclude that the resolution and precision of the present dataset allows the detection of preconception changes that may correlate with lifestyle behaviors.

### Complications of pregnancy show distinct dynamical changes

So far, we have considered healthy pregnancies. To study complications of pregnancy, we analyzed data from pregnancies diagnosed with three major complications – pre-eclampsia (5,629 pregnancies, 1.80%), gestational diabetes (7,233 pregnancies, 2.31%) and postpartum hemorrhage (4,566 pregnancies, 1.46%). The incidence of these complications in the dataset is lower than expected(*29*–*31*). The lower incidence can be attributed to exclusion of risk factors, such as chronic illness (**Methods**) and maternal age over 35 years (*30, 32, 33*). We compared the test dynamics to healthy pregnancies during preconception, gestation and postpartum **(Fig. 5)**.

**Fig. 5.**
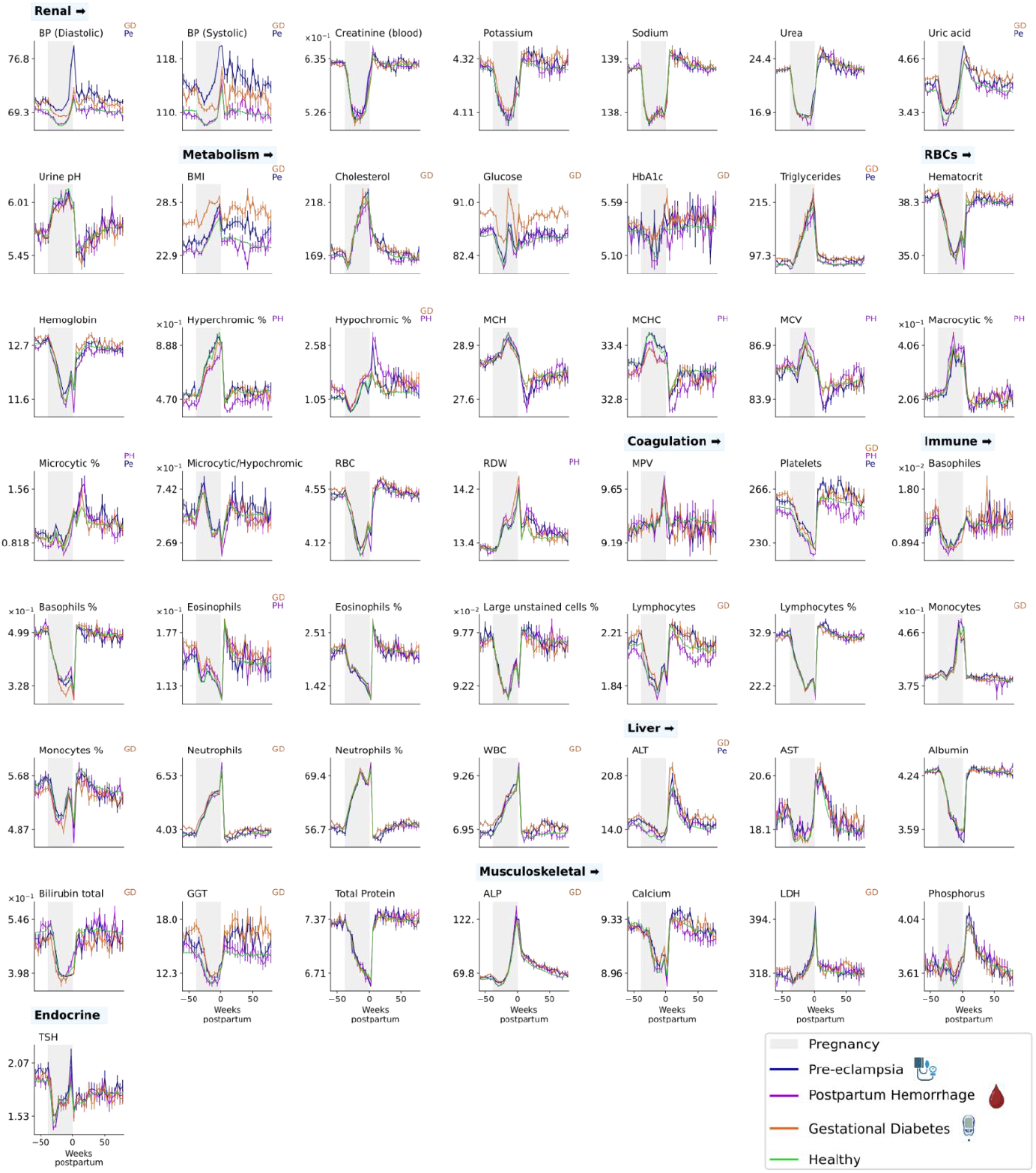
Dynamics of lab tests in complications of pregnancy. Tests with significant differences from healthy pregnancy (green) are marked by *Pe* for pre-eclampsia (blue), *PH* for postpartum hemorrhage (purple), and *GD* for gestational diabetes (orange). The y-axis is different between tests, for units see **table S1**. For the same figure with quantile scores, see **fig. S5**.

Pre-eclampsia is a complex disorder of pregnancy characterized by high blood pressure, headaches and visual disturbances(*34*). In some cases (<2% of preeclamptic patients(*35*)), pre-eclampsia develops into eclampsia, a life-threatening condition usually requiring urgent delivery(*36*).

The causes of pre-eclampsia are not fully understood; it is believed to involve factors related to the placenta, the immune system, and genetics (*32, 37*). We find that 7 tests deviated significantly from the healthy reference (**Fig. 6A,D,G,J**). These include elevated platelets and ALT in the preconception period, elevated gestational uric acid, elevated postpartum triglycerides and high systolic and diastolic blood pressure throughout the study period. High blood pressure during pregnancy is the main diagnostic tool for pre-eclampsia coupled with diagnosis of proteinuria (*37*)

**Fig. 6:**
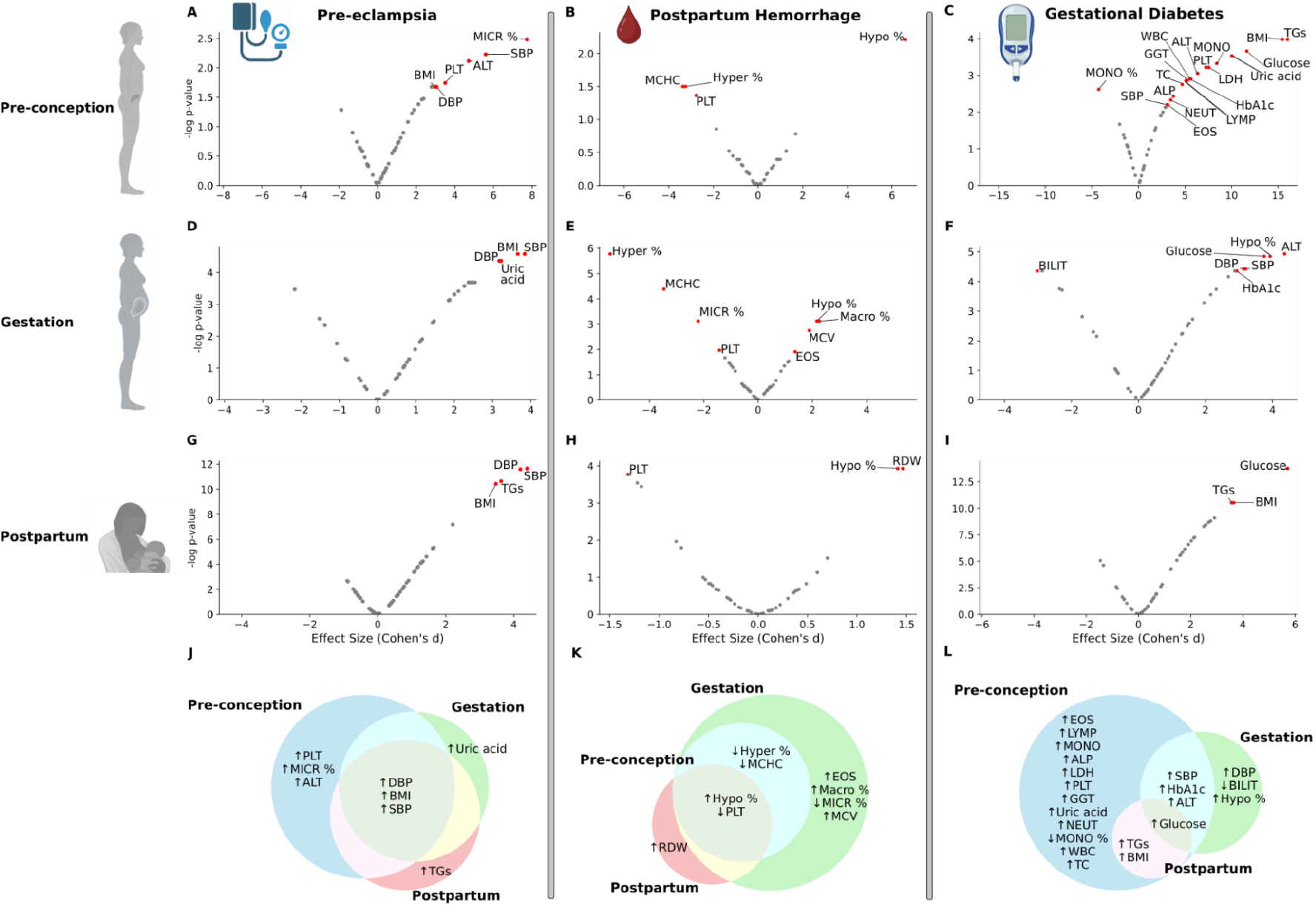
Tests with significant deviations from healthy pregnancies in pre-eclampsia, gestational diabetes and postpartum hemorrhage. **A-I**) volcano plots of -log p-value (FDR corrected) versus effect size for each test at each time point. Significant tests are marked with a red dot and their name (Methods). **J-L**) Venn diagrams showing the significant tests in the preconception (blue), gestation (green) and postpartum (red) periods for each complication.

Postpartum hemorrhage (PPH) is a major cause of maternal morbidity and mortality characterized by excessive bleeding (≥1000ml) after childbirth, typically within 24 hours after delivery and up to 12 weeks postpartum (*38*–*40*). The primary cause is uterine atony, where the uterus fails to contract adequately after delivery(*40*). Other causes include retained placental fragments, tears in the cervix or vaginal tissues, and coagulation disorders(*40*). We find that 9 tests deviate significantly from the healthy reference, including tests before delivery **(Fig. 6B,E,H,K)**. Platelets are mildly reduced, suggesting altered blood clotting even before pregnancy. Other coagulation markers are not significantly altered(*41, 42*). PPH is also associated with a distinctive pattern of decreased MCHC (Mean Corpuscular Hemoglobin Concentration) and elevated MCV (Mean Corpuscular Volume) before conception and during gestation. This agrees with a longitudinal study where a higher MCV and MCH towards the end of pregnancy was associated with higher likelihood of PPH(*43*) **(Fig. 6K)**.

Gestational diabetes is characterized by high blood glucose that develops during pregnancy in females who did not previously have diabetes(*44*). It usually appears in the second or third trimester and can affect the health of both mother and fetus(*45, 46*). We find that 20 tests deviate significantly from the healthy reference **(Fig. 6C,F,I,L)**. This includes high glucose and HbA1c, elevated GGT liver damage test, and elevated triglycerides before and after pregnancy. These values are associated with obesity and inflammation, both causes of insulin resistance which contributes to gestational diabetes.

In all three complications, some of the significant changes are seen before conception or after delivery rather than during gestation (**Fig. 6J,K,L**). In gestational diabetes, 17 of the 20 significantly different tests are different during preconception, of these 12 are statistically different solely in the preconception period. In other words, during gestation the dynamical profiles are generally similar to healthy pregnancies. This is interesting given that tests for diagnosing pathologies such as GD and pre-eclampsia are done during gestation(*47, 48*).

## Discussion

We present a cross-sectional dataset of 40 million lab test measurements from 300,000 pregnancies during a 140-week period spanning preconception, gestation and postpartum. The dataset is unprecedented in terms of number of participants and time intervals and covers all major laboratory tests. About half of the tests take on the order of months to a year to return to baseline after delivery, highlighting the physiological aftermath of pregnancy. During gestation all tests show sizable changes, and about half show large overshoots after delivery. The precision of the dataset allows detection of intricate dynamical changes, including the impact of preconception supplements, and the deviations from healthy pregnancy in pre-eclampsia, gestational diabetes and postpartum hemorrhage. This study thus provides a resource for understanding pregnancy and the postpartum period and demonstrates how it may be used to understand mechanisms in systems physiology.

This study greatly expands our knowledge of the postpartum period, since most postpartum studies considered only one or a few timepoints. Rather than a ‘fourth trimester’ with rapid return to baseline, there is a slow recovery time of between 10 and 50 weeks for 32/76 of the tests. Examples of such slowly adapting tests are alkaline phosphatase, albumin, AST and ALT as well as sodium and uric acid.

We find that the postpartum return of the tests to baseline occurs by a trajectory that differs from the trajectory of change during pregnancy - a phenomenon called hysteresis. Postpartum adaptation is a distinct physiological process and not merely the reverse of pregnancy dynamics.

Several tests show a difference between their preconception values and their values 80 weeks postpartum. These postpartum differences include elevated levels of the inflammation marker CRP and reduced mean cell hemoglobin MCH and iron. The differences could result from postpartum behavioral changes and/or from lasting physiological effects of pregnancy. Telling these factors apart is a major question for future research.

The lab tests show two types of stereotypical profiles - either a smooth rise-and-fall, where delivery redirects the direction of change back to baseline, or jump-like, where delivery causes a sharp overshoot or undershoot. Rebounds and sharp reversals have not been systematically characterized previously because studying them requires many temporal intervals which were lacking in most previous studies.

These profile shapes can be rationalized based on general physiological principles. Overshoots are consistent with a compensatory mechanism that grows during pregnancy and remains high after delivery causing overcompensation. An example of such compensation occurs in the thyroid axis, where thyroid functional mass grows during pregnancy under control of TSH and hCG, increasing the capacity to produce thyroid hormones. This extra mass takes months to recover postpartum given the slow turnover of thyroid cells, causing overshoot dynamics in thyroid hormones. The ability of endocrine glands to change mass has important beneficial functions, such as dynamic compensation of variation in physiological parameters (*49*). Gland mass changes add a timescale of months to hormone dynamics and contribute to hormone seasonality (*50*), explain subclinical endocrine diseases (*17*) and cause extended dysregulation after chronic stress is relieved (*51*).

Pathologies of pregnancy showed distinct temporal profiles in specific tests. These differences from healthy pregnancies were more pronounced before conception and after delivery than during gestation for many of the tests. Several aberrations were shared between two pathologies - gestational diabetes and pre-eclampsia - suggesting the possibility of a pan-complication signature.

This study presents detailed cross-sectional temporal trajectories of pregnancy and postpartum physiology. These trajectories reveal prolonged recovery times and overshoot effects of many tests after delivery, preconception dynamics of many tests, and perturbed tests in pregnancy complications at unprecedented detail. It suggests processes that allow the mother’s physiology to adapt to the multisystemic load of pregnancy and to navigate the abrupt effects of delivery. The power of this dataset stems from having delivery as a well-defined temporal signpost, a t=0. A similar approach might be useful for understanding other temporal transitions such as growth and development in childhood, puberty, menopause, and the course of specific diseases (diagnosed at t=0) and their recovery processes. We hope that the present dataset will lead to a better understanding of pregnancy and postpartum biology and inspire similar studies of other crucial physiological processes that unfold over time.

## Limitations of the study

This study has limitations associated with use of medical datasets, including the effects of ascertainment bias. This study considered pregnancies in a single country; future work can consider effects of different locales. The study is cross-sectional and should be tested by future longitudinal studies that can assess subtypes of pregnancy trajectories.

## Acknowledgments

We thank all members and Dr. Yaniv Ovadia of our lab for discussions. We thank Gabi Barabash and Ran Balicer for the Clalit−Weizmann collaboration. Data acquisition was approved by the Clalit Helsinki Committee RMC-1059-20. Figure 1A and figure 6 were made with BioRender (BioRender.com).

## Author contributions

Conceptualization: AB, YKK, YT, UA

Methodology: AB, NMC, AM, RM, YKK, UA

Formal Analysis: AB, RM, AM, NMC

Visualization: AB, RM

Writing – Original Draft: AB, UA

Writing – Review and Editing: RM, AB, UA

## Data and materials availability

The source code and data used to perform the analysis is available at the GitHub repository as of the date of publication. Github repository:

https://github.com/AlonLabWIS/PregnancyMillions

