## Supplementary Materials for "Pregnancy and postpartum dynamics revealed by millions of lab tests"

Alon Bar *et al.*

**This PDF file includes:**

Methods
Supplementary Text
Figs. S1 to S5
Tables S1 to S3

### Methods

Study Population

The study population consisted of individuals from the Clalit healthcare database(*11*, *52*). We considered all pregnancies of females aged 20 to 35 between 2003 and 2020. Information about pregnancies before 2002 is not available. We estimated the fraction of first pregnancies for the years 2010-2020 to reduce the influence of first pregnancies before 2003 which we cannot account for. For more information, see "stats.csv" in the GitHub repository.

Data Collection

Medical records were pseudonymized by hashing of personal identifiers and randomization of dates by a random number of weeks uniformly sampled between 0 and 13 weeks for each patient and adding it to all dates in the patient diagnoses, laboratory, and medication records. This randomization does not affect timing relative to delivery.

We examined the timeframe of 60 weeks before delivery to 80 weeks after delivery for all documented labors within our study population. We identified deliveries by ICD9 code V27 and confirmed a childbirth record for the individual. We excluded preterm deliveries (<37 weeks, ICD9 code 644) stillbirths and labors with more than one newborn. Nonetheless, 12% of deliveries were found to be <37 weeks and missing the 644 code.

To mitigate ascertainment bias of the test results, for each test, we removed data from individuals with chronic disease that affects the test if the onset of the disease was up to 6 months after the test. We also removed data from individuals who purchased drugs which affected the tests in the 6 months before the tests. Chronic diseases are defined as non-pediatric ICD9 codes with a Kaplan−Meyer survival drop of >10% over 5 years. An exhaustive list of chronic disease can be found under `non_healthy_icd_and_cohort.csv` in the GitHub repository. Drugs that affect a test were defined as drugs with significant effect on the test (false discovery rate < 0.01). This step allowed us to focus on a relatively healthy subset of the pregnant population, reducing the confounding effects associated with specific health conditions listed above or medication usage(*12*).

To exclude the potential effect of follow-up pregnancies in the 80 weeks following delivery, we excluded lab values from individuals with another delivery within 60 weeks following the measurement.

For each pregnancy, we gathered all available test values including standard blood count, kidney and liver function tests, blood coagulation tests, lipid panel, inflammation markers and hormones (**table S1**). We then discretized test values into time points relative to the time of birth in weekly intervals for each test. In addition to test values, we also extracted data on patients including age (at measurement, mean and interquartile range) and BMI (the most proximal BMI measurement in medical records outside pregnancy, mean and interquartile range, if available).

Quantile transformation

We transformed each measurement into a quantile score, normalized between 0 and 1. Quantiles were computed from cumulative distributions of test values from a reference population of age-matched non-pregnant females ($F_{test,age}$). The non-pregnant reference population included healthy non-medicated females according to the R package LabNorm(*12*), including individuals in the study cohort during the time periods that they were not pregnant. The transformation is summarized as:

$$\text{quantile\_score} = F_{test,age}(\text{value})$$

Data aggregation

Each individual test result included the age, latest BMI (if given), week postpartum (with 0 being delivery), and the quantile score. We aggregated the data into summary statistics: For each weekly interval per lab test, we calculated the mean, standard deviation and (5, 10, 25, 50, 75, 90, 95) percentiles of all the above-mentioned data types. To obtain a test value at the mean quantile score for figures 1 and 5, we performed a back-transformation by transforming mean quantiles into test level values. The back transformation uses a sparsely sampled cumulative distribution function per test (denoted $F^{-1}$ below) and linear interpolation between the quantiles. The values for $F^{-1}$ were queried using LabNorm and can be found in the attached git repo in a file named "Labnorm.csv". Tests without such reference were filtered (see **Data filtering** below). Age parameterizing the back-transform is the median age of the population at the same week (week $i$ below) as the mean quantile back-transformed:

$$mean\_value_i = {F^{-1}}_{test,age_i}(mean\_quantile\_score_i)$$

Error bars are quantile standard error of the mean (SEM) likewise back-transformed to test value **(table S2)**. Fluctuations between neighboring points suggest additional errors on the order of 40-180% of SEM.

Data filtering

The original dataset included 110 tests. Tests with too few measurements, either for a specific gestational week or averaging on all weeks, were excluded. The procedure resulted in excluding 31 tests with a low number of test results. If the standard error of the mean at any weekly interval, or the mean across all intervals, exceeded a threshold the test was deemed noisy. 3 additional tests were discarded with this procedure. Tests with no reference for the back transformation did not have enough test results, therefore excluded in advance. For filtered tests and thresholds see **table S2,3**.

Data analysis

Unless otherwise stated, the mean quantile values were used for each test and weekly interval. See **Quantile transformation** and **Data aggregation** above for more details.

Averaging consecutive time points (smoothing)

In **Fig. 1C** we smoothed the curves for tests with smaller sample sizes by using averaged consecutive weekly intervals. This process was performed for visualization and was not used for data analysis. See **fig. S3** for the data without smoothing.

Dynamic range

We assessed the dynamic range across pregnancy by comparing the time-intervals with the minimum and maximum values for each test. We performed a two-sample t-test for each test between these time-intervals with correction for false discovery rate based on the Benjamini–Hochberg method with a threshold of 0.05 using FDR correction in the Statsmodels Python package.

Clustering

We performed clustering to group together tests with similar profiles. Tests were clustered (ward) using a distance metric of $1 - r_s$ where $r_s$ is the Spearman correlation and `fclust` from the python module `scipy.clustering.hierarchy`. Each test is a vector of the quantile score at each of the 140 week-intervals. For more information see **fig. S4**.

Principal component analysis

We performed PCA using the 'PCA' implementation in the python package sklearn .

Settling time

We define the postpartum baseline using the quantile values in the last 10 weeks of the dataset, weeks 70-80 after delivery, and define standard deviations using the average of the standard deviations of test values in these 10 bins. We next smoothed the mean test value data using a Gaussian kernel smoother, using the gaussian_filter function of the Python scipy.ndimage with sigma=1 and mode='nearest'. The settling time is the time after which at least 90% of the smoothed timepoints remain within 0.2 standard deviations of the baseline values. The cutoff of 0.2 standard deviations was chosen following a visual inspection of the data. In **Fig. 3** settling time was computed from quantile scores.

Preconception dynamics

We used linear regression to model the relationship between the test mean quantile score and time (in weeks) in the preconception period (60 to 38 weeks before delivery). We considered a test to have a significant dynamical trend if the absolute value of the regression line slope was greater than 0.1 and the p-value was less than 0.05 after controlling for false discovery rate (FDR). Linear regression was performed using 'curve_fit' from python Scipy 'optimize'.

Data Processing (pregnancy complications)

The methodology mentioned above was used to create a dataset of pregnancies with diagnosis of pre-eclampsia, gestational diabetes and postpartum hemorrhage using the ICD9 codes 642.4-9 excluding 642.8, (648.83, 648.81) excluding 250 (diabetes) and 666 respectively. The measurement was included if the diagnosis was made not more than 140 weeks before/after the measurement was taken, and if the diagnosis was made in the [-42, +3] week where 0 denotes delivery. The test results were aggregated at a 4-week interval due to a lower number of measurements. Filtering as mentioned above yielded a subset of 50 tests, for a list see **table S1**.

Data analysis (pregnancy complications)

Test results for each condition (Pre-eclampsia, Gestational Diabetes, Postpartum hemorrhage) were compared against the unaffected dataset aggregated at a 4-week interval. We performed a paired t-test and separated the weekly intervals to preconception, gestation and postpartum. We corrected for multiple comparisons using the Benjamini–Hochberg method with a threshold of <0.05, and Cohen's d as the effect size for a paired t-test:

$$Cohen's\ d = \frac{t\_statistic}{\sqrt{n}}$$

Where "$n$" is the number of data points per period. We set effect size thresholds corresponding to large effect sizes(*53*, *54*) by visual inspection, as follows: Pre-eclampsia ≥ 3.0, postpartum hemorrhage ≥ 1.3, gestational diabetes ≥ 3.0.

**Supplementary Text**

Minimal mathematical model of dynamical profiles **(Fig. 2K)**

We consider a simple mathematical model based on the incoherent feed-forward loop (IFFL) circuit (*15*) with input $u(t)$ - designating the physiological load of pregnancy- and variables $X$ and $Y$. In this model, factor $X$, a first order system, is produced in proportion to the input $u(t)$ at rate $a$ and removed at rate $b$:

$$1)\quad dX/dt = au(t) - bX$$

The second factor in this model, $Y$, stands for a test for which there is a compensation process, namely $X$. Similar to $X$, $Y$ is produced in proportion to the input $u(t)$ at rate $c$ and removed at rate $d$. However, $Y$ has an added layer of regulation where the compensation mechanism $X$ reduces $Y$ production rate:

$$2)\quad \frac{dY}{dt} = c\frac{u(t)}{X} - d\,Y$$

We simulated the model over 140 weeks starting with $u(t) = 1$ for 22 weeks (preconception), followed by a pregnancy load of 38 weeks where $u(t)$ rises linearly from 1 with a slope of $m = 0.01/week$, and returns to 1 postpartum. Note that here $t = 0$ is conception and not delivery:

$$3)\quad u(t) = \begin{cases} 1, & t \leq 0 \\ 1 + m \cdot t, & 0 \leq t < 38 \\ 1, & t > 36 \end{cases}$$

Removal rates were $b = 1/20\ weeks$ and $d = 1/\ weeks$ to provide a slow timescale for $X$, and a fast time scale for $Y$. Other parameters were chosen to give a steady state of 1. To solve the equations, we will assume that $Y$ dynamics are faster than $X$ and faster than changes in the input dynamics $\mathrm{u(t)}$ . Where $t = 0$ is conception and $t = 38$ is delivery. Solving $Y$ at quasi steady state $\left(\frac{\mathrm{dY}}{\mathrm{dt}}\ = 0\right)$:

$$4)\quad \mathrm{Y_{qst}}\ =\ \frac{\mathrm{c\,u(t)}}{\mathrm{d\,X(t)}}$$

The steady state of the model is $\left(\frac{\mathrm{dY}}{\mathrm{dt}}\ = 0,\ \frac{\mathrm{dX}}{\mathrm{dt}}\ = 0,\ \mathrm{u(t) = u_0}\right)$:

$$5)\quad \mathrm{X_{st}}\ =\ \frac{\mathrm{a}}{\mathrm{b}}\mathrm{u_0}$$

$$6)\quad \mathrm{Y_{st}}\ =\ \frac{\mathrm{c\,b}}{\mathrm{d\,a}}$$

We modeled the pregnancy load as input that rises with gestational time:

$$7)\quad \mathrm{u(t) = (1 + mt)}$$

The time-dependent solution of $X$ for the pregnancy period is given by solving the $X$ differential equation with time varying input (Eq. 1, 3).

$$8)\quad \mathrm{X(t)} = \frac{\mathrm{a}}{\mathrm{b}}\mathrm{(1 + mt)} + \frac{\mathrm{a\,m}}{\mathrm{b^2}}\left(\mathrm{e^{-bt}} - 1\right)$$

This function is asymptotically linear in $t$ (considering pregnancy even for $t > 38$):

$$9)\quad X(t_{\rightarrow\infty}) = \frac{a}{b^2} mt + O(1)$$

To find the pregnancy time-dependent solution of $Y$, we substitute $X(t)$ (Eq. 8) and $u(t)$ (Eq. 7) to $Y_{qst}$ (Eq. 6):

$$10)\ Y(t) = \frac{c(1+mt)}{d\left(\frac{a}{b}(1+mt)+\frac{a}{b^2}\left(m+me^{-bt}\right)\right)}$$

This function is asymptotically constant:

$$11)\ Y(t_{\rightarrow\infty}) = \frac{c\,b}{d\,a} + O(1) = Y_{st}$$

At delivery, following 38 weeks of gestation (Eq. 7,10, $t = 38$):

$$12)\ X_{delivery} = \frac{38am}{b} + \frac{b_X}{b} + \frac{b_X m}{b^2} + \frac{a \cdot me^{-38 \cdot b}}{b^2}$$

$$13)\ Y_{delivery} = \frac{c(1+38m)}{d\left(\frac{38am}{b}+\frac{a}{b}+\frac{am}{b^2}+\frac{ame^{-a_X 38}}{b^2}\right)}$$

Postpartum, the pregnancy load is removed and $u(t) = 1$. $X$ decays to its original steady state:

$$14)\ X(t) = \frac{a}{b} + \frac{\left(bX_{delivery} - a\right)e^{-bt}}{b}$$

And $Y$ follows (substituting Eq. 14 in Eq. 5):

$$15)\ Y(t) = \frac{c}{d\left(\frac{a}{b}+\frac{\left(bX_{delivery}-a\right)e^{-bt}}{b}\right)}$$

<u>Calculating noise threshold</u>

See **table S2** for notation and **table S3** for thresholds.

We define $\sigma_{F_x^{-1}}$ to be the standard deviation of the back transformed values:

$$\sigma_{F_x^{-1}} = \sqrt{\mathbb{E}\left[\left(\left[\widetilde{\mu_{x_\iota}}\right] - \mathbb{E}_i\left[\widetilde{\mu_{x_\iota}}\right]\right)^2\right]} = \sqrt{\frac{1}{140-1}\sum_{i=-60}^{80}\left(\widetilde{\mu_{x_\iota}} - \overline{\widetilde{\mu_{x_\iota}}}\right)^2}$$

We define $\sigma_{i_{gF_x^{-1}}}^{err}$ to be the standard error of the mean at week $i$ for test $x$ for the back transformed values:

$$\sigma_{i_{F_x^{-1}}}^{err} = \frac{1}{2} \cdot \left(F_x^{-1}\left(\min\left\{\mu_{x_i}^q + \frac{\sigma_{x_i}^q}{\sqrt{n_{x_i}}}, 1\right\}, \lfloor\alpha_{x_i}\rfloor\right) + F_x^{-1}\left(\max\left\{\mu_{x_i}^q - \frac{\sigma_{x_i}^q}{\sqrt{n_{x_i}}}, 0\right\}\right), \lfloor\alpha_{x_i}\rfloor\right)$$

* For the “complications” dataset there are a total of 35 time point over the same 140 weeks period (a 4-week resolution).

System-by-System overview of pregnancy and postpartum dynamics

We provide a system-by-system overview of the physiological dynamics observed in the dataset. The dataset captures the major trends with much finer detail than previously known. It reveals previously unknown temporal changes such long recovery times postpartum, and overshoots and rebound effects in nearly all physiological systems.

**Kidney**

Renal physiology is affected by pregnancy in multiple ways. There is an increase in renal volume, blood flow and glomerular filtration rate (GFR), as well as an increase in reabsorption of nutrients and electrolytes(55). After delivery, diuresis causes elimination of excess fluids within days(7) and GFR returns to normal within 6-8 weeks (56). Kidney volume can take up to 6 months to decrease to pre-pregnant volume(57).

The kidney removes waste products from the blood including creatinine, urea and uric acid(58). The dataset shows that the corresponding test values drop sharply in the first trimester, consistent with the temporal profile of increased GFR(55, 59, 60) . Urea and creatinine remain low until delivery, whereas uric acid rises in the last trimester, peaking at delivery, and normalizes over months postpartum. Blood electrolytes (Na, Cl, K, Mg) show a drop at the first trimester and remain low until delivery, consistent with the GFR profile. The observed patterns of electrolytes are in line with the known decrease in serum osmolarity(55, 61, 62). We find that most of these tests show a previously unreported undershoot or overshoot postpartum.

Urine composition, controlled by the kidneys, is also subject to multiple changes. The dataset shows that urinary pH becomes slightly more alkaline during pregnancy (63). It shows a pattern that is inverse to other renal blood tests such as creatinine, urea, and sodium, including an undershoot postpartum. Specific gravity, which reflects urine particle concentration, shows a gradual decline during pregnancy (64)11, which is resolved within several weeks postpartum. **(fig. S2A,B)**

**Liver**

The liver changes to support the growing fetus and the metabolic demands of the mother's body. Placental hormones, such as estrogen and progesterone, affect hepatic metabolic, synthetic and excretory functions (*65*).

The liver produces most of the blood proteins including albumin and globulin (*66*). The dataset shows that albumin and total protein concentration tests drop gradually during the first trimester and stay low until delivery and then recover over a few weeks postpartum. This follows the temporal profile of maternal blood volume and may be attributed to uncompensated hemodilution(*67*). In contrast, globulin stays relatively constant. Bilirubin tests report on the ability of the liver to clear bilirubin. They show a U-shaped curve, unlike previously reported(*68*), with a nadir around mid-pregnancy.

Liver enzyme tests GGT, ALT and AST are used to detect liver damage. In the dataset, their dynamics differ - GGT shows a drop in pregnancy like bilirubin and rapid recovery, whereas ALT and AST do not drop as much during pregnancy but show a pronounced overshoot after delivery. This is surprising since AST and ALT are usually considered unchanged in pregnancy(*65, 67*). The overshoot in ALT levels is similarly unreported thus far(*65*) **(fig. S2C,D)**.

**Musculoskeletal system**

Weight gain and the growing fetus impose forces and biochemical stress on the maternal skeleton, muscles and ligaments (*3*, *12*). Demand for calcium and phosphorus rises during pregnancy and lactation, and bone metabolism is affected by estrogen deficiency during lactation. Bone mass recovery can take up to a year after delivery(*3*).

The dataset shows indicators of muscle and bone damage or turnover, including alkaline phosphatase (ALP), lactate dehydrogenase (LDH) and creatinine kinase (CK), changing strongly over pregnancy and postpartum. ALP and LDH drop in the first trimester and then rise to peak around delivery (*62*). Postpartum, the tests return to baseline, with LDH normalizing faster than ALP. The rise in ALP is thought to be due to the production of placental and bone isoenzymes(*65*). CK tests also drop in the first trimester and rise again towards delivery(*13*, *69*). Postpartum CK begins normal, undershoots and adapts to baseline within weeks.

The dataset shows that calcium and phosphorus, major bone components, differ in their dynamics. Calcium drops during pregnancy, with a nadir around week 20, probably due to increased fetal demand (*70*). It shows a jump following delivery and rises to the normal range within weeks. Calcium levels then slowly rebound during postpartum. In contrast to the strong changes in calcium, phosphorus dynamics are complex but generally close to the normal range, with a postpartum overshoot that slowly rebounds. Negative calcium balance is seen several weeks postpartum and the prolonged increase in serum phosphorus might be a result of adaptation mechanisms that support the calcium demand of lactation (*70*) **(fig. S2E)**.

**Red blood cells**

During pregnancy, red blood cell concentration (RBC), volume fraction (HCT) and hemoglobin (Hgb), tests which reflect oxygen carrying capacity, all show the same U-shape decline in the dataset that approaches the lower bound of the normal range. This is due to hemodilution that is only partially compensated by increased RBC production leading to dilutional anemia(*71*). These tests show sharp changes around delivery, possibly due to hemorrhage, and return to pre-pregnancy levels within weeks. **(fig. S2F)**

Mean cell volume (MCV), mean cell hemoglobin (MCH) rise during pregnancy. Mean cell hemoglobin concentration (MCHC) also rises in this period, indicating that the increase in hemoglobin content is not exclusively due to an increase in cell volume. These variables show a postpartum undershoot. For about 20 weeks postpartum, MCH decreases more than MCV, in agreement with the observed undershoot in MCHC. Red cell size distribution width RDW rises and then adapts to normal levels with an additional peak postpartum. These effects are consistent with increased RBC production(*72*) and iron-deficiency anemia(*73*). **(fig. S2F)**

Ferritin, which reflects the body's iron stores, shows a gradual decline with a nadir during the third trimester. During postpartum it overshoots for about 20 weeks, parallel to an undershoot in MCHC. Blood iron shows a mild rise during the first trimester, but then declines until delivery. Hemoglobin, ferritin and iron show a sequence of nadir timing during mid-late pregnancy, potentially reflecting a hierarchy in iron metabolism. Transferrin, a protein that transports iron in the circulation, increases dramatically during pregnancy(*74*). It shows a rebound as it decreases and undershoots postpartum. Taken together, these test dynamics reflect the increased demand for iron and the adaptation processes that increase its bioavailability(*75*)**(fig. S2G).** Additional RBC-related tests are shown in **fig. S2H**.

**Immune system**

The maternal immune system changes to protect the mother and fetus from pathogens and to provide tolerance to the allogeneic fetus (*76*) **(fig. S2I-K)**. Lymphocytes in the dataset show U-shaped dynamics during pregnancy, with a nadir around mid-pregnancy and an overshoot postpartum. Neutrophils, the most abundant white blood cells, and total white blood cell counts (WBC) both rise and plateau in the third trimester (*76*). They return to normal range within a few weeks postpartum, but then undershoot and take about 30 weeks to reach stable value. Monocytes show a mild rise during pregnancy(*76*) and return to baseline shortly after delivery. Eosinophil and basophil counts, which are generally thought to be unchanged in pregnancy(*76*) are shown in the dataset to decrease mildly during pregnancy and overshoot postpartum. **Fig. S2**I shows the blood counts, and **fig. S2J** the relative counts.

The inflammatory marker CRP (C-reactive protein) increases during pregnancy(*76*), peaking shortly after delivery. The dataset shows that CRP remains slightly higher than pre-pregnancy baseline for up to 80 weeks. The apparent above-baseline levels could be explained by the lower-than-expected levels of CRP observed before pregnancy (see **Health style behaviors are reflected in preconception dynamics**) Procalcitonin, a marker of bacterial infection, decreases during pregnancy and normalizes rapidly after delivery. A previous study showed an opposite trend(*77*). Large unstained cells, a marker for viral infection, show a mild decrease during pregnancy **(fig. S2K)**.

**<u>Coagulation</u>**

Coagulation undergoes multiple changes and generally shifts towards a hypercoagulable state(*78*). Fibrinogen, a procoagulant factor, rises during pregnancy(*78*), and undershoots postpartum **(fig. S2L,M)**. It reaches normal range within a few weeks but takes about 20 weeks to stabilize. During pregnancy clotting becomes more rapid(*78*, *79*) as seen in the decline in clotting tests APTT-R, APTT-sec and PT-INR, which resolves within a few weeks postpartum. Platelet count in the dataset shows a decrease due to hemodilution(*80*), with a slight overshoot postpartum. We also observe changes in mean platelet distribution width (PDW), which increases during pregnancy, indicating changes in production and removal that counter hemodilution.

**<u>Metabolism</u>**

Metabolism changes to ensure nutrient supply for the fetus. These changes are regulated primarily by human placental lactogen (hPL) secreted by the placenta(*81*).

The dataset in **fig. S2O** shows that HbA1c, a test used to assay glucose levels averaged over 2-3 months, drops in the first trimester and then rises, mirroring the decline and rise of insulin resistance(*82*, *83*).

Postpartum, the hPL-dependent insulin resistance that developed during pregnancy is relieved rapidly, but compensation by insulin-secreting beta cells is still strong and lasts for weeks(*49*, *82*, *84*). This may be an example of a rebound effect due to a compensatory mechanism, potentially explaining the drop in glucose in the first weeks of postpartum.

In early pregnancy, the maternal body begins to store fat, and serum lipids increase(*81*, *85*), as seen in the dataset by a rise in cholesterol and triglycerides that slows down towards delivery. Postpartum, many of these tests show a rapid small drop, perhaps due loss of hPL signals, and then a slower recovery over months. The prolonged recovery of these tests is possibly due to the mother's adipose tissue stores that decrease over weeks, and breastfeeding that affects lipids metabolism and body mass(*86*, *87*) **(fig. S2N-P).**

**Fig. S1.**

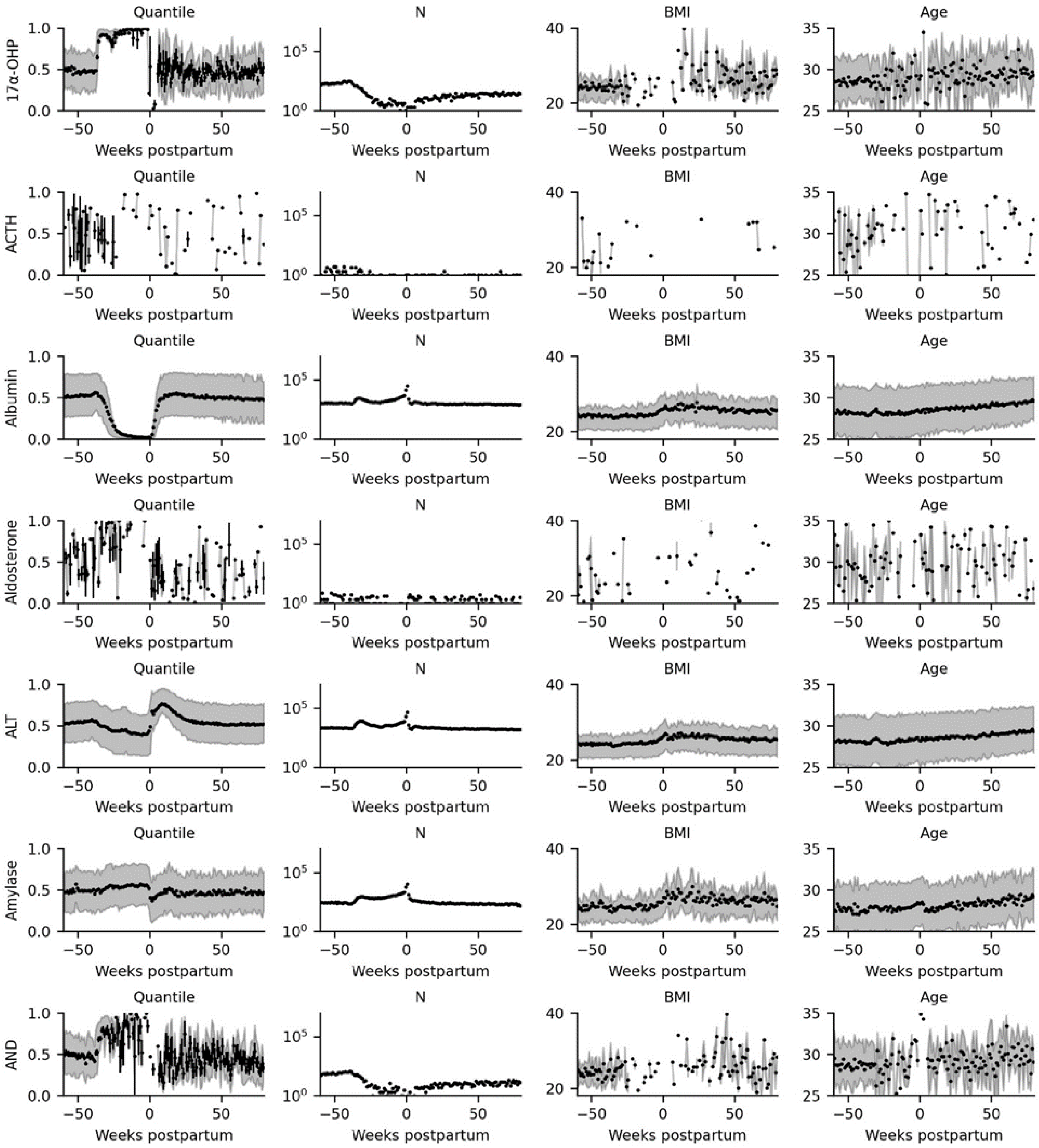
Quantile
N
BMI
Age
17α-OHP
ACTH
Albumin
Aldosterone
ALT
Amylase
AND
Weeks postpartum

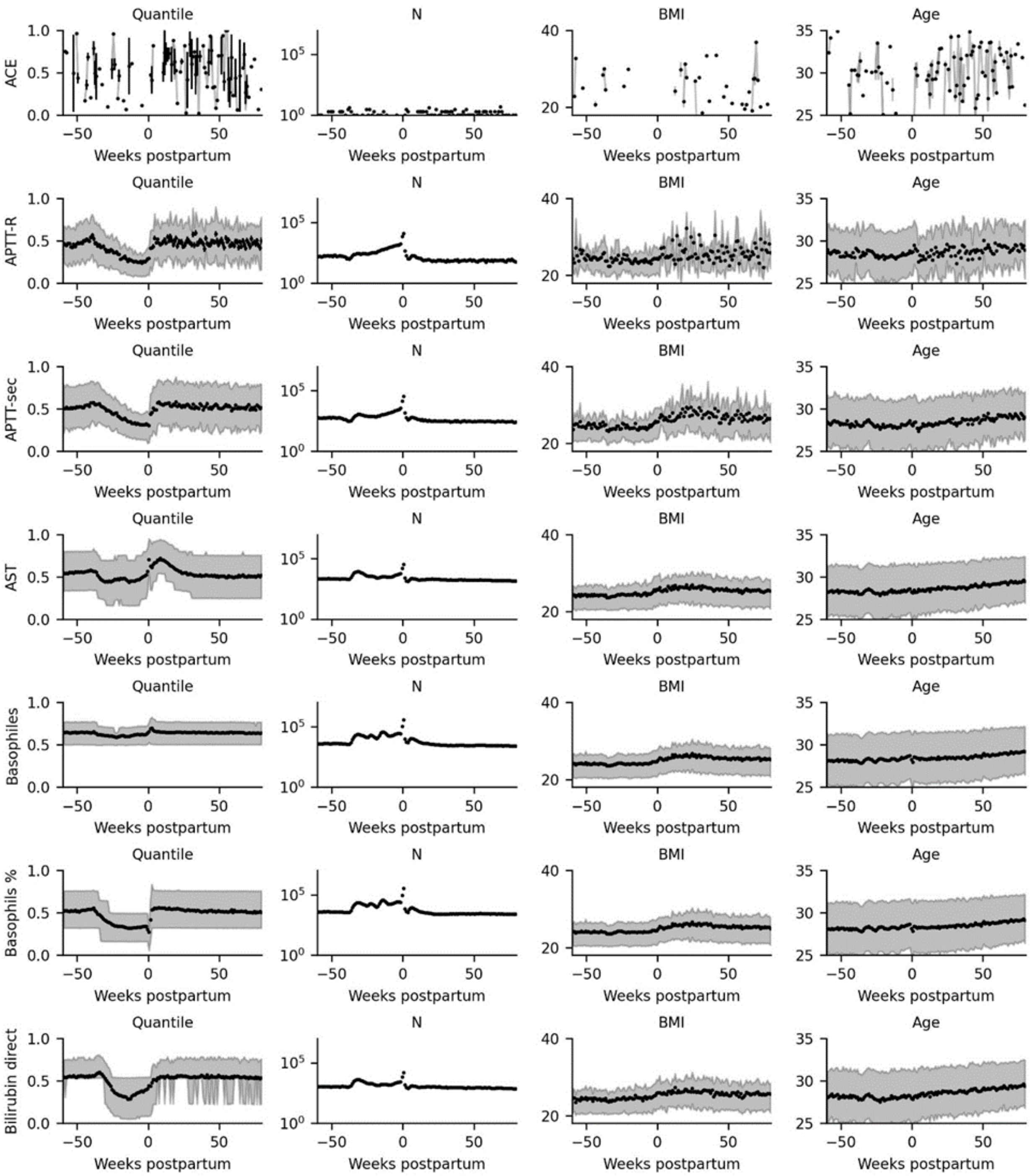
Quantile
N
BMI
Age
ACE
APTT-R
APTT-sec
AST
Basophiles
Basophils %
Bilirubin direct
Weeks postpartum

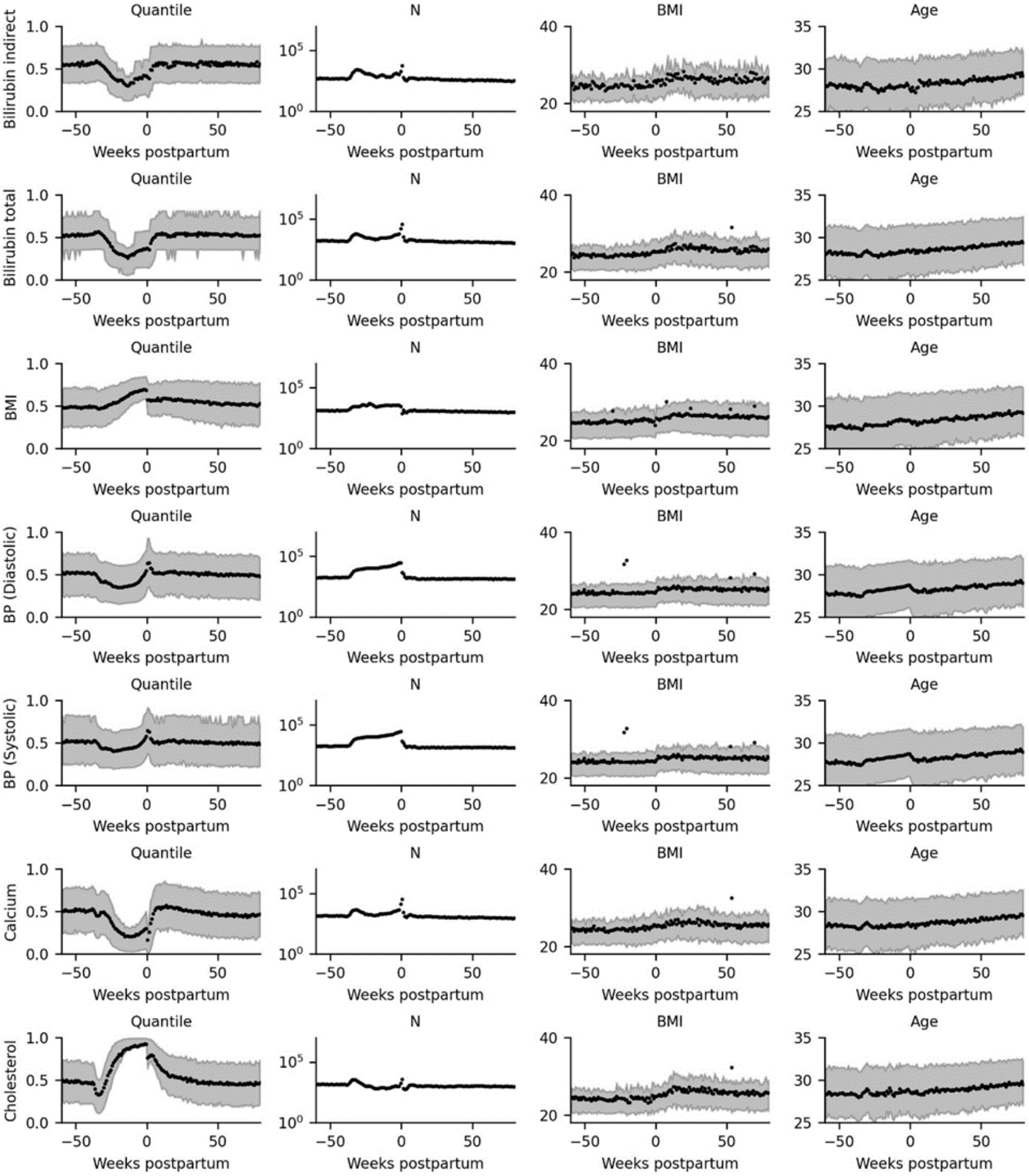
Quantile
N
BMI
Age
Bilirubin indirect
Bilirubin total
BMI
BP (Diastolic)
BP (Systolic)
Calcium
Cholesterol
Weeks postpartum

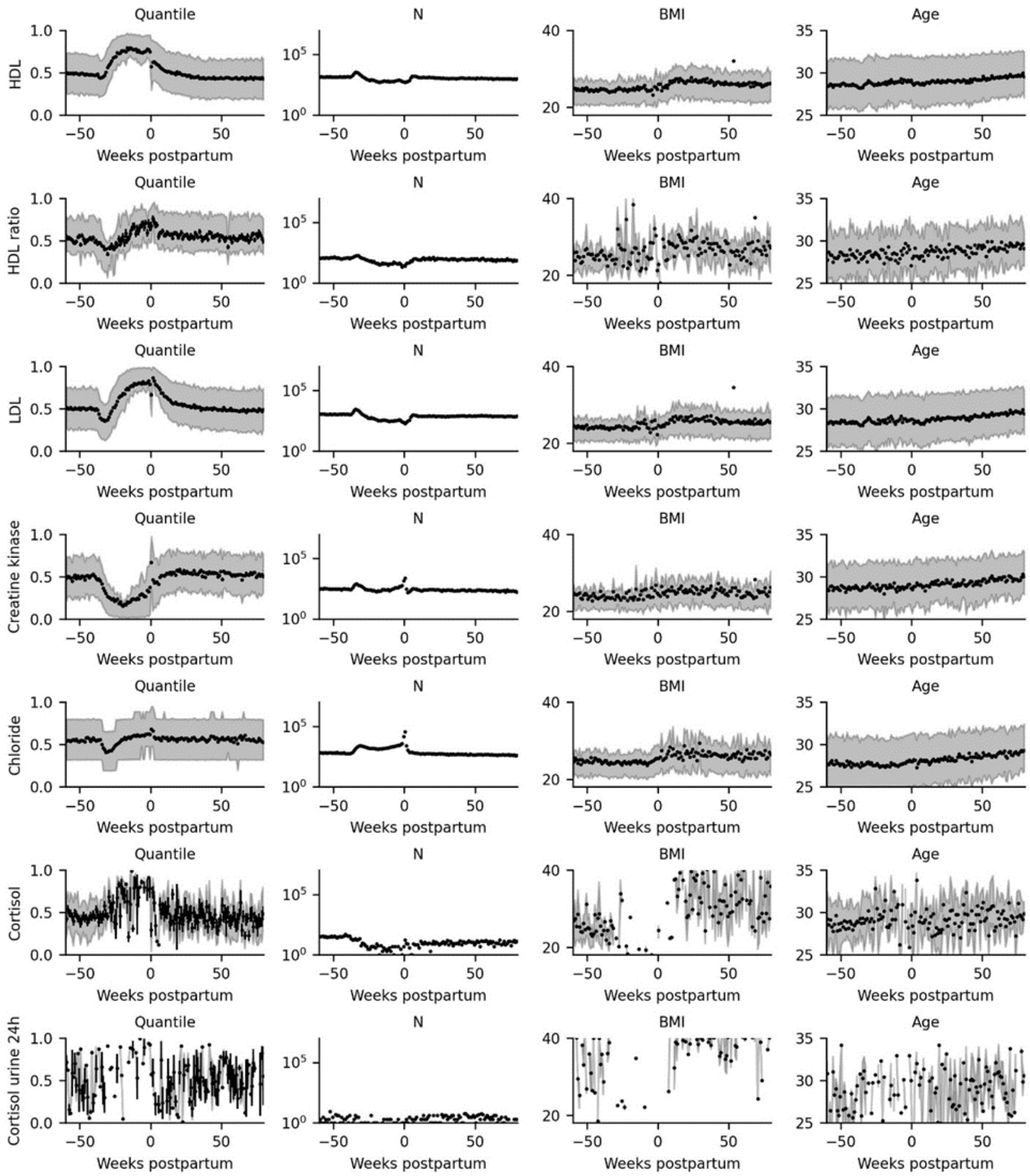

Quantile
N
BMI
Age
HDL
HDL ratio
LDL
Creatine kinase
Chloride
Cortisol
Cortisol urine 24h
Weeks postpartum

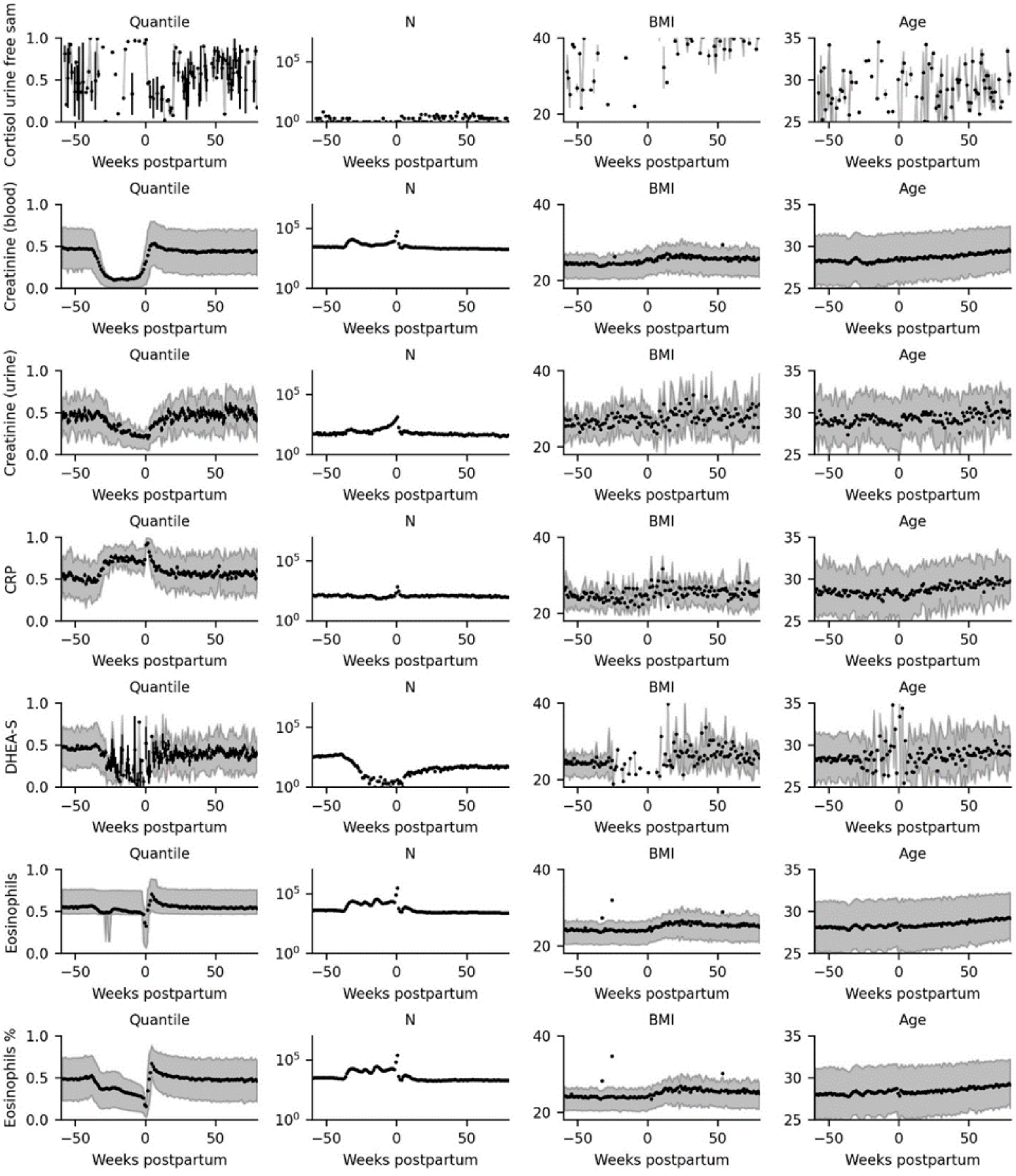

Quantile
N
BMI
Age
Cortisol urine free sam
Creatinine (blood)
Creatinine (urine)
CRP
DHEA-S
Eosinophils
Eosinophils %
Weeks postpartum

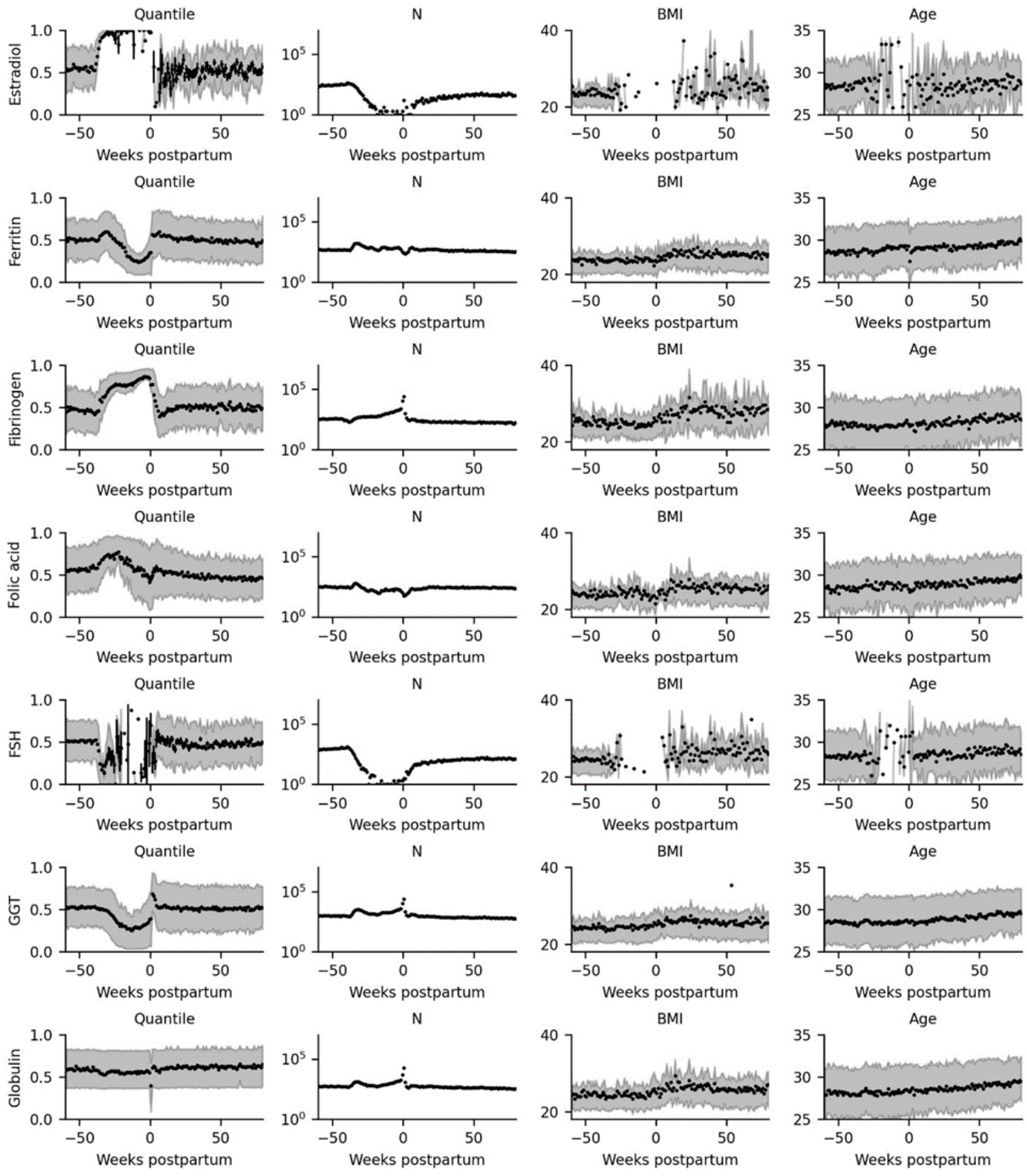

Quantile
N
BMI
Age
Estradiol
Ferritin
Fibrinogen
Folic acid
FSH
GGT
Globulin
Weeks postpartum

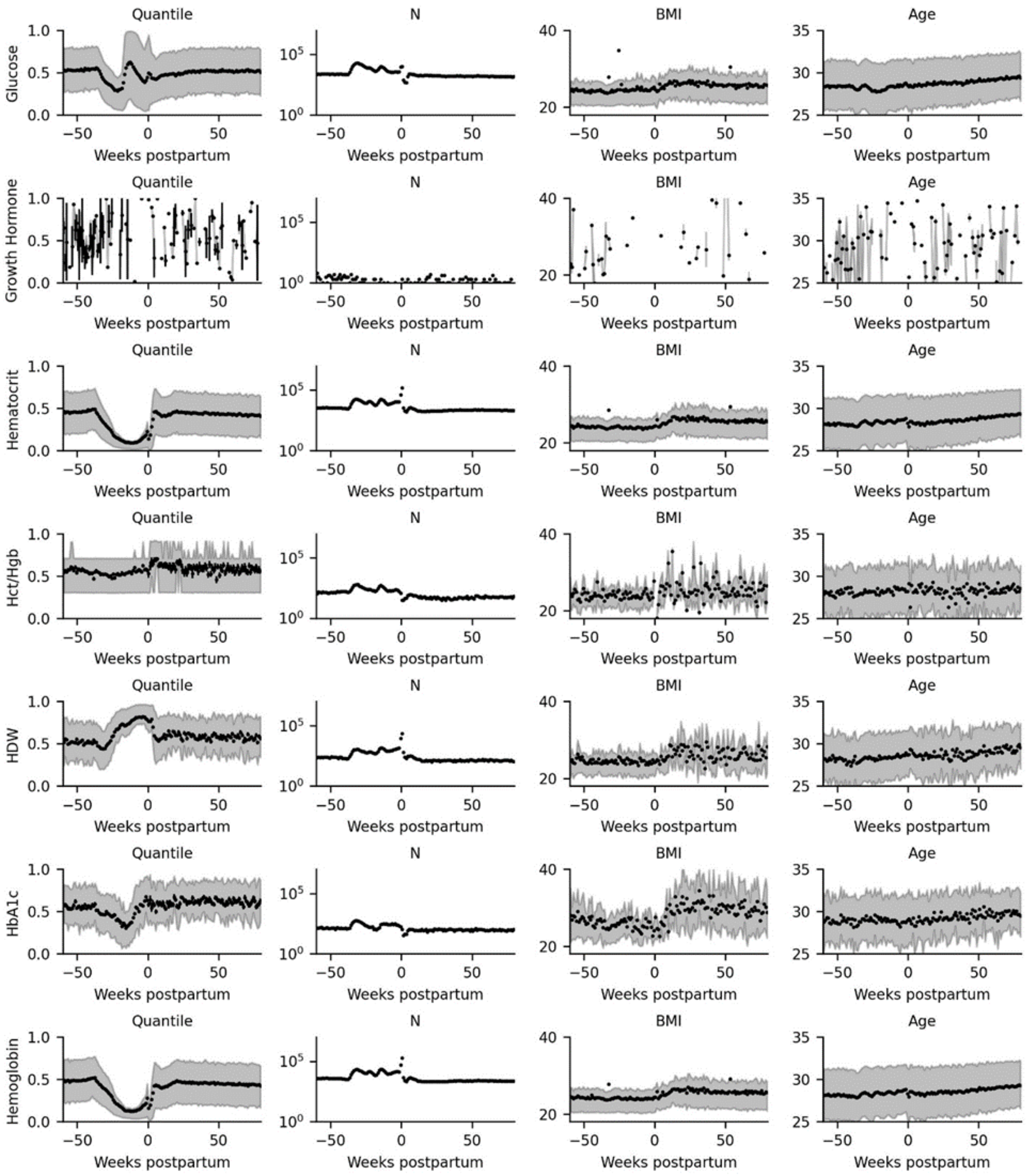
Quantile
N
BMI
Age
Glucose
Growth Hormone
Hematocrit
Hct/Hgb
HDW
HbA1c
Hemoglobin
Weeks postpartum

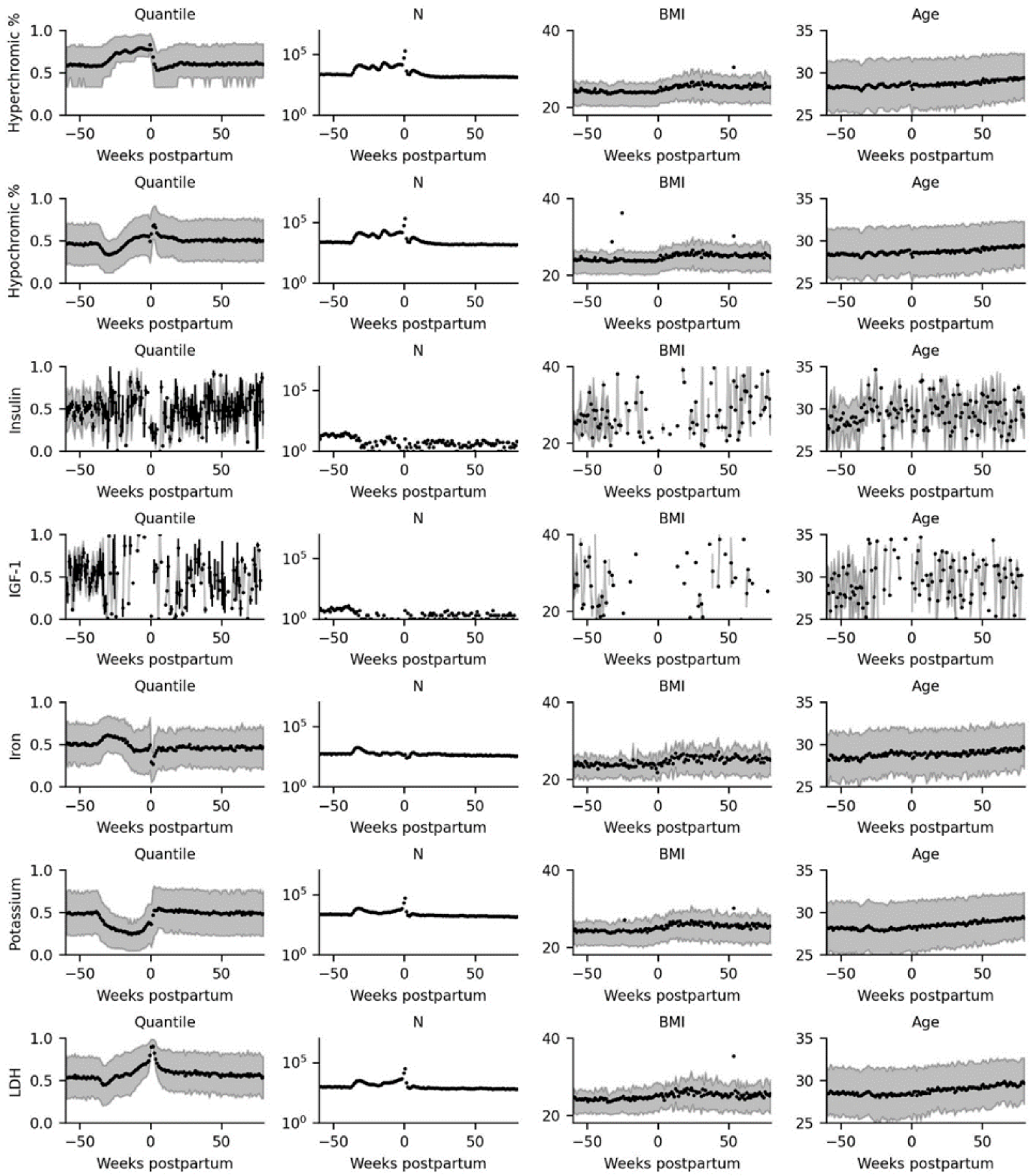

Quantile
N
BMI
Age
Hyperchromic %
Hypochromic %
Insulin
IGF-1
Iron
Potassium
LDH
Weeks postpartum

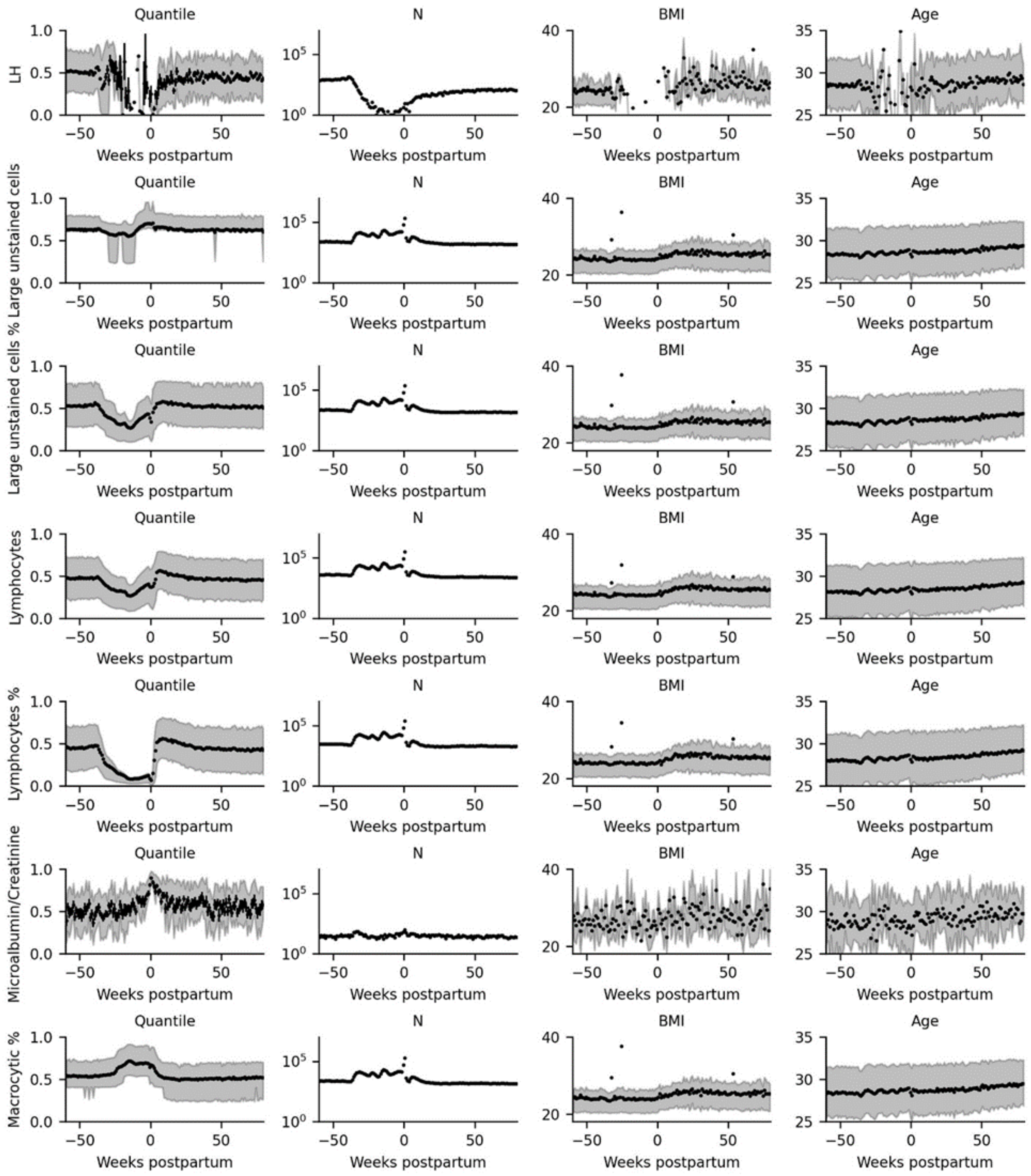
Quantile
N
BMI
Age
LH
Large unstained cells
Large unstained cells %
Lymphocytes
Lymphocytes %
Microalbumin/Creatinine
Macrocytic %
Weeks postpartum

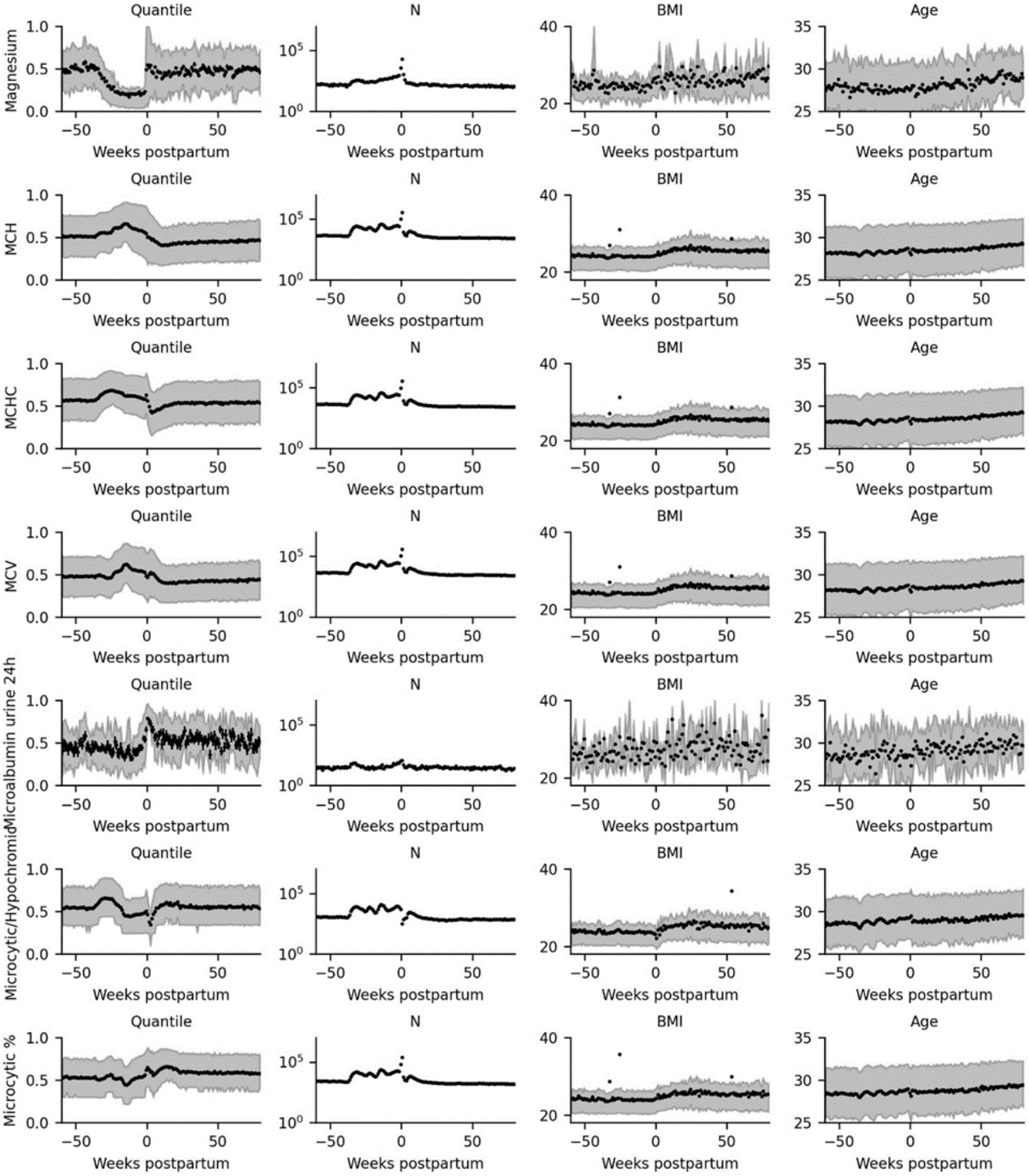
Magnesium
MCH
MCHC
MCV
Microalbumin urine 24h
Microcytic/Hypochromic
Microcytic %
Quantile
N
BMI
Age
Weeks postpartum

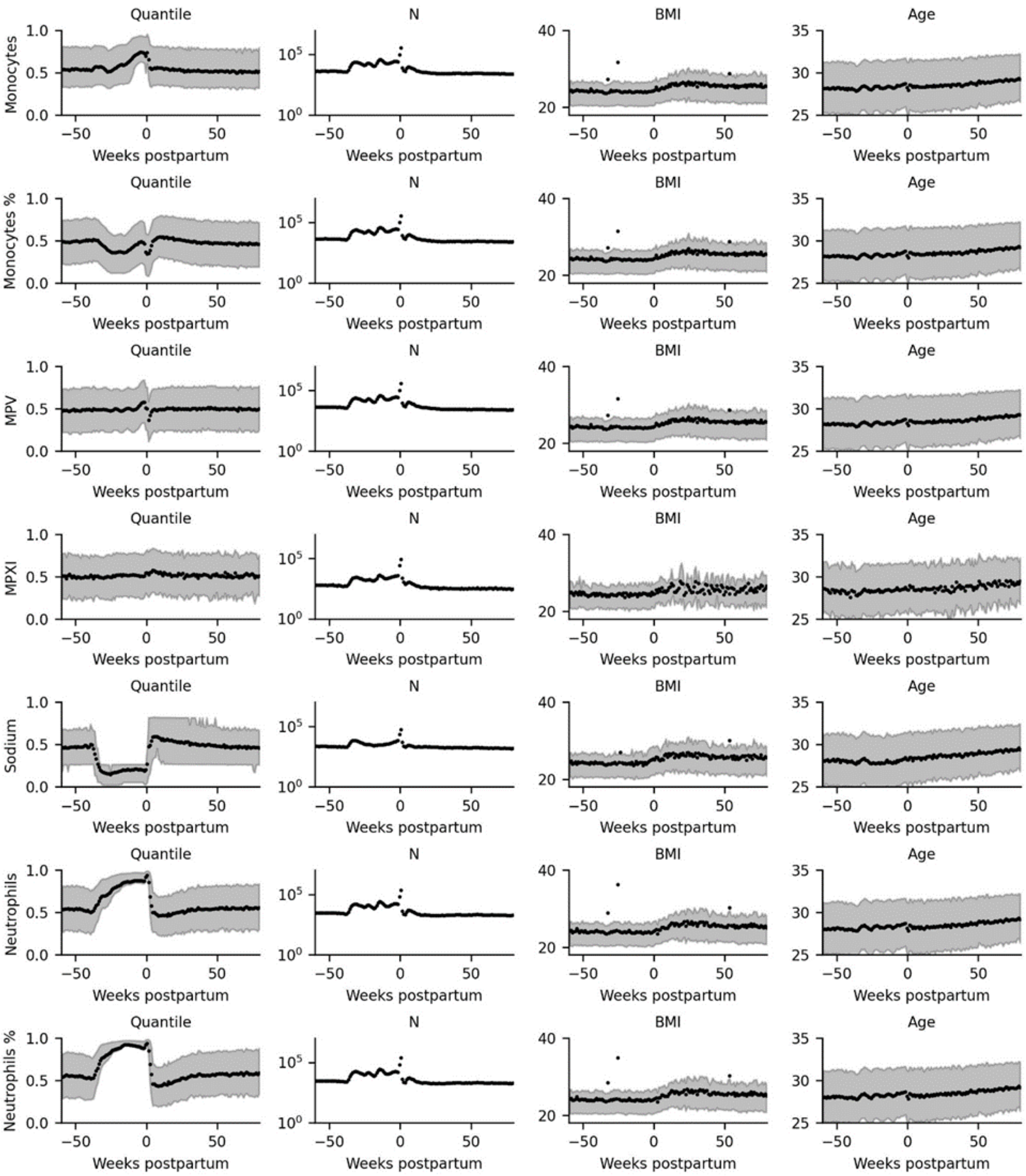
Quantile
N
BMI
Age
Monocytes
Monocytes %
MPV
MPXI
Sodium
Neutrophils
Neutrophils %
Weeks postpartum

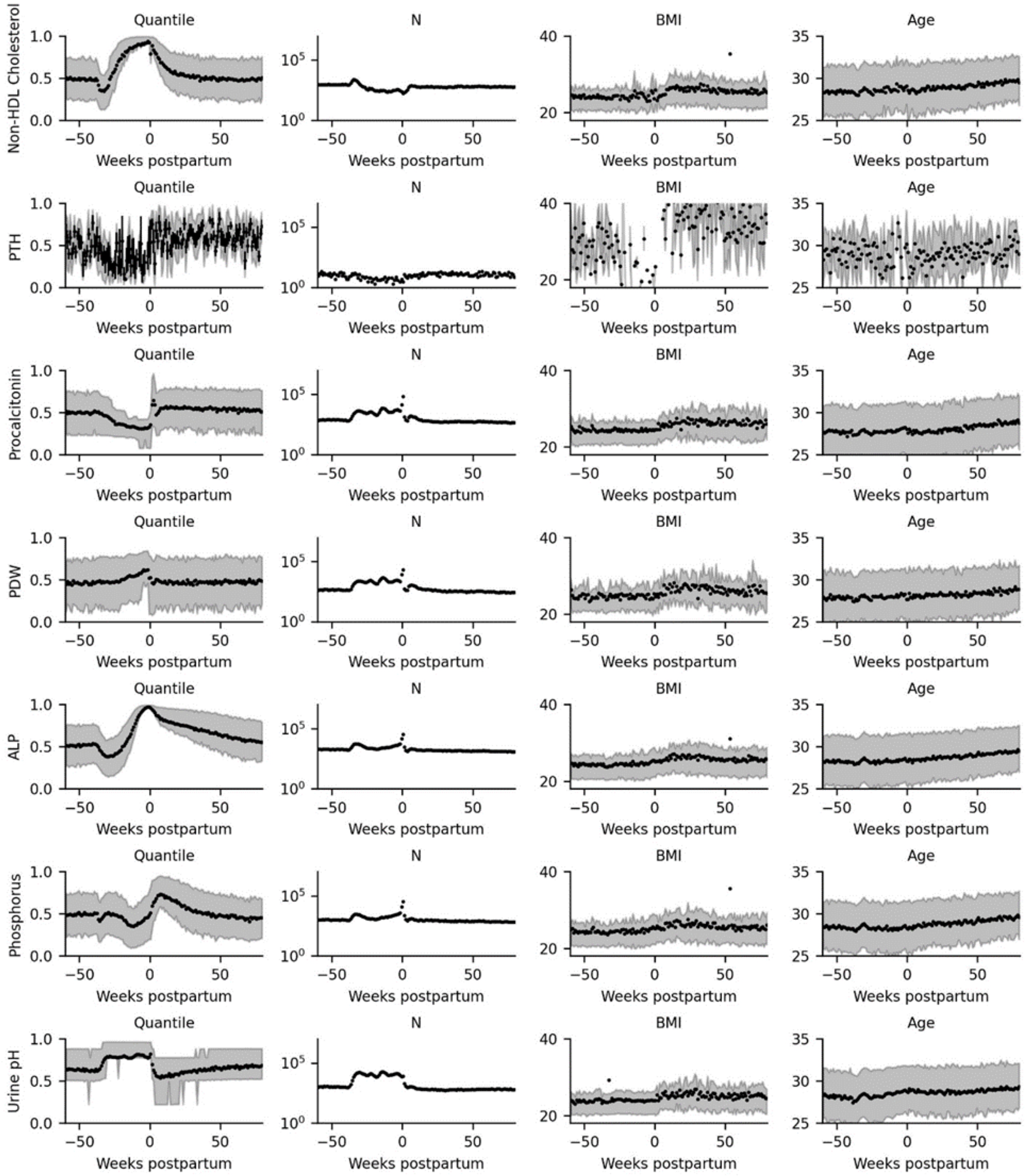
Non-HDL Cholesterol
PTH
Procalcitonin
PDW
ALP
Phosphorus
Urine pH
Quantile
N
BMI
Age
Weeks postpartum

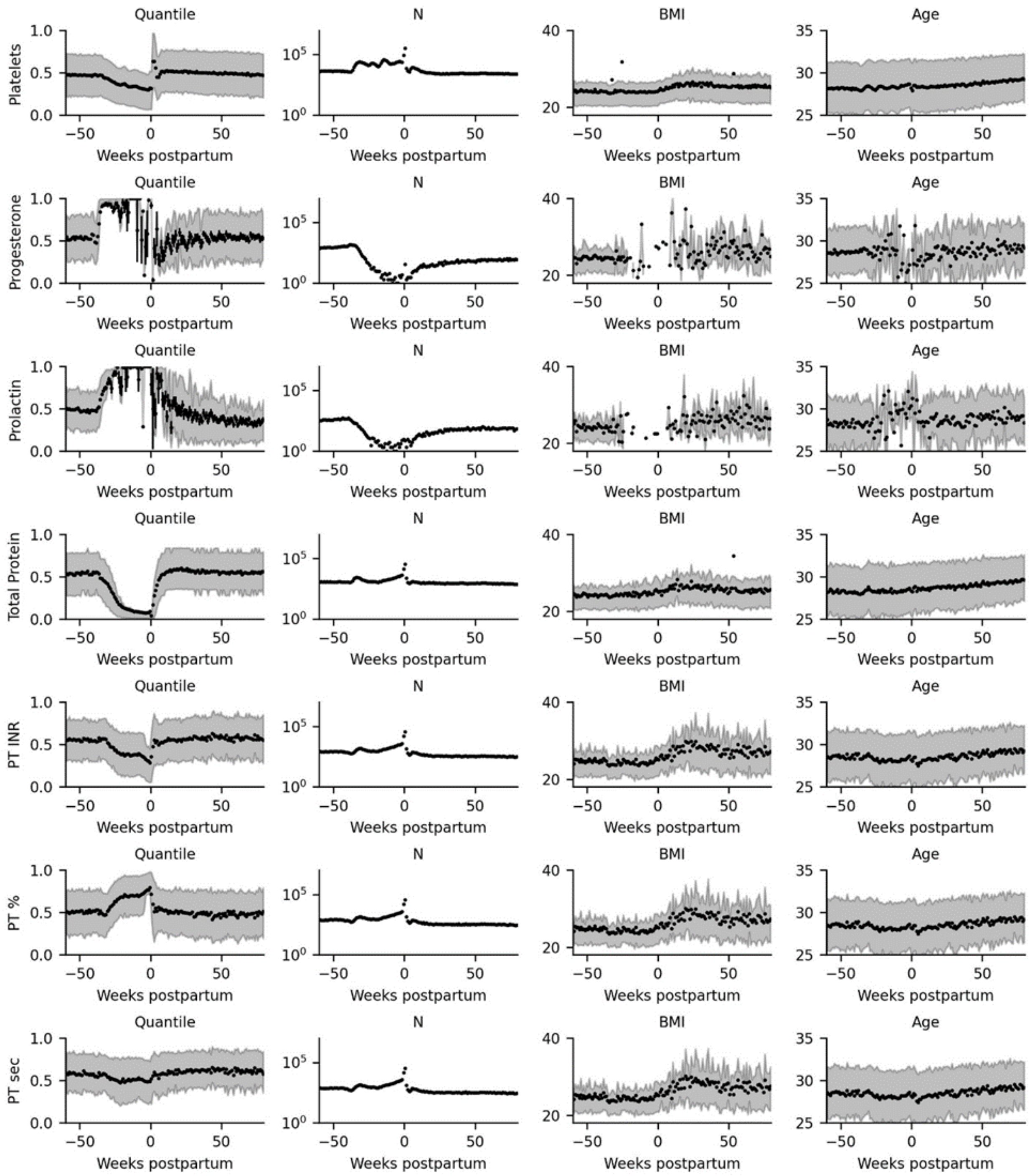

Quantile
N
BMI
Age
Platelets
Progesterone
Prolactin
Total Protein
PT INR
PT %
PT sec
Weeks postpartum

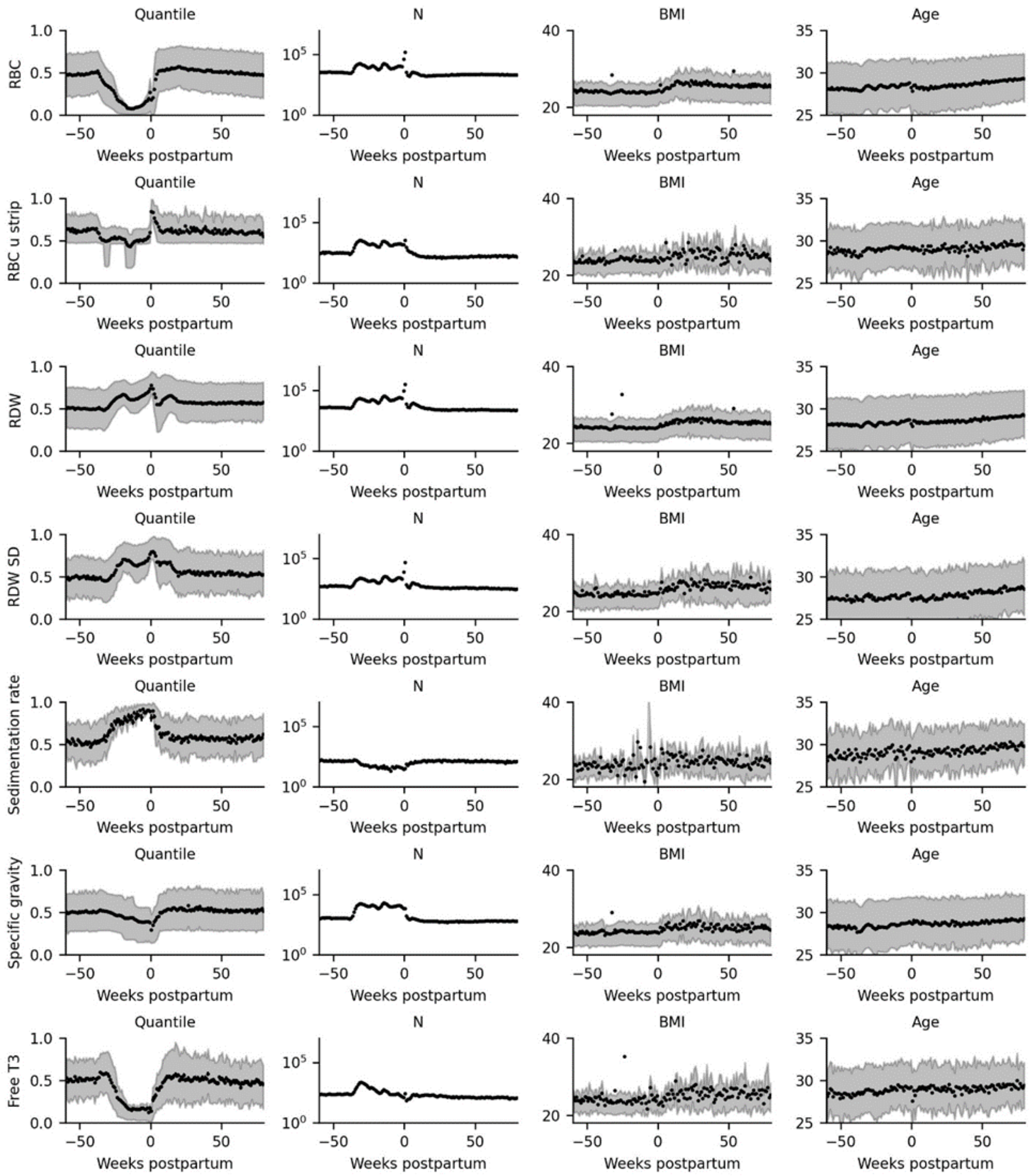

Quantile
N
BMI
Age
RBC
RBC u strip
RDW
RDW SD
Sedimentation rate
Specific gravity
Free T3
Weeks postpartum

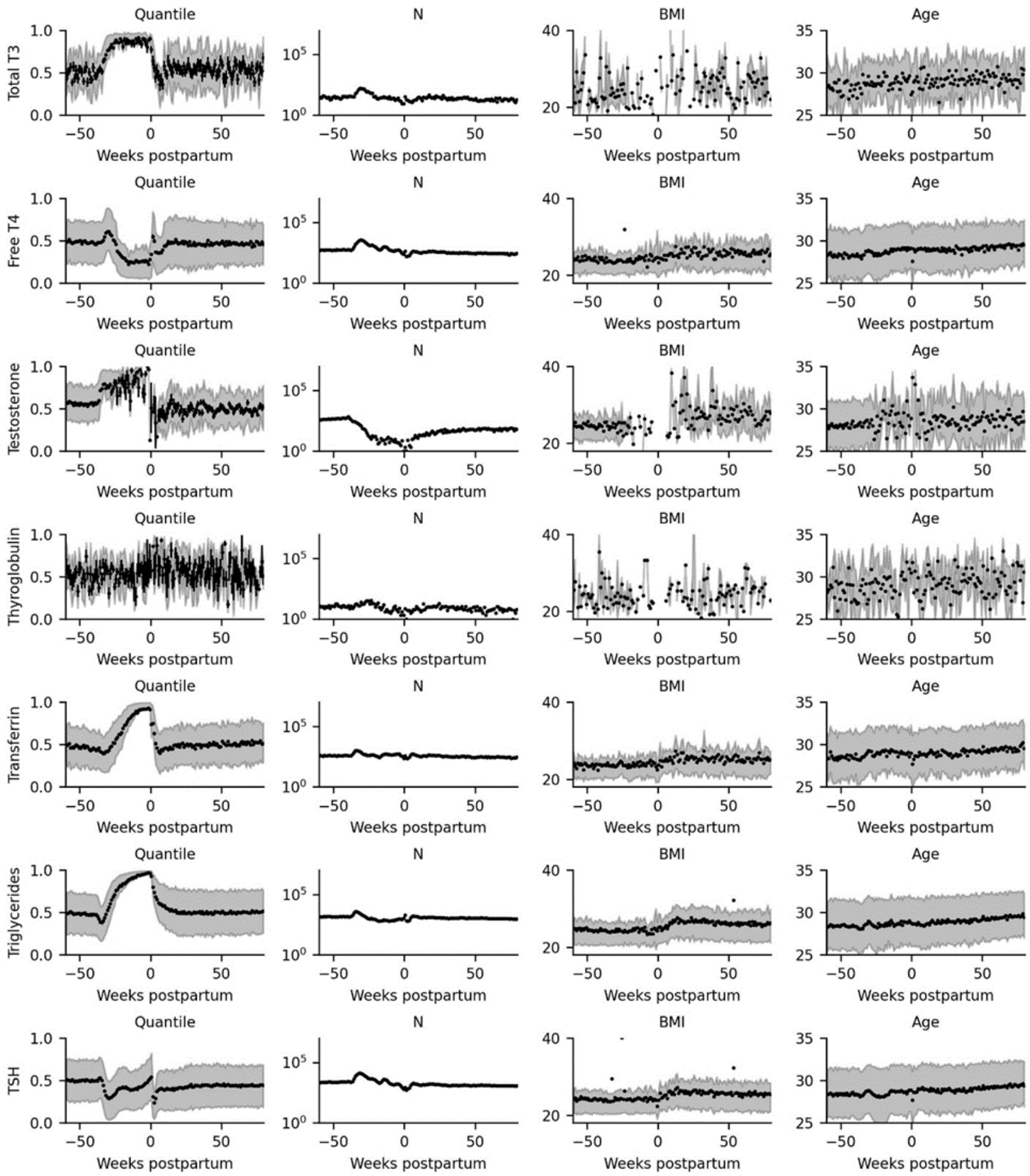
Quantile
N
BMI
Age
Total T3
Free T4
Testosterone
Thyroglobulin
Transferrin
Triglycerides
TSH
Weeks postpartum

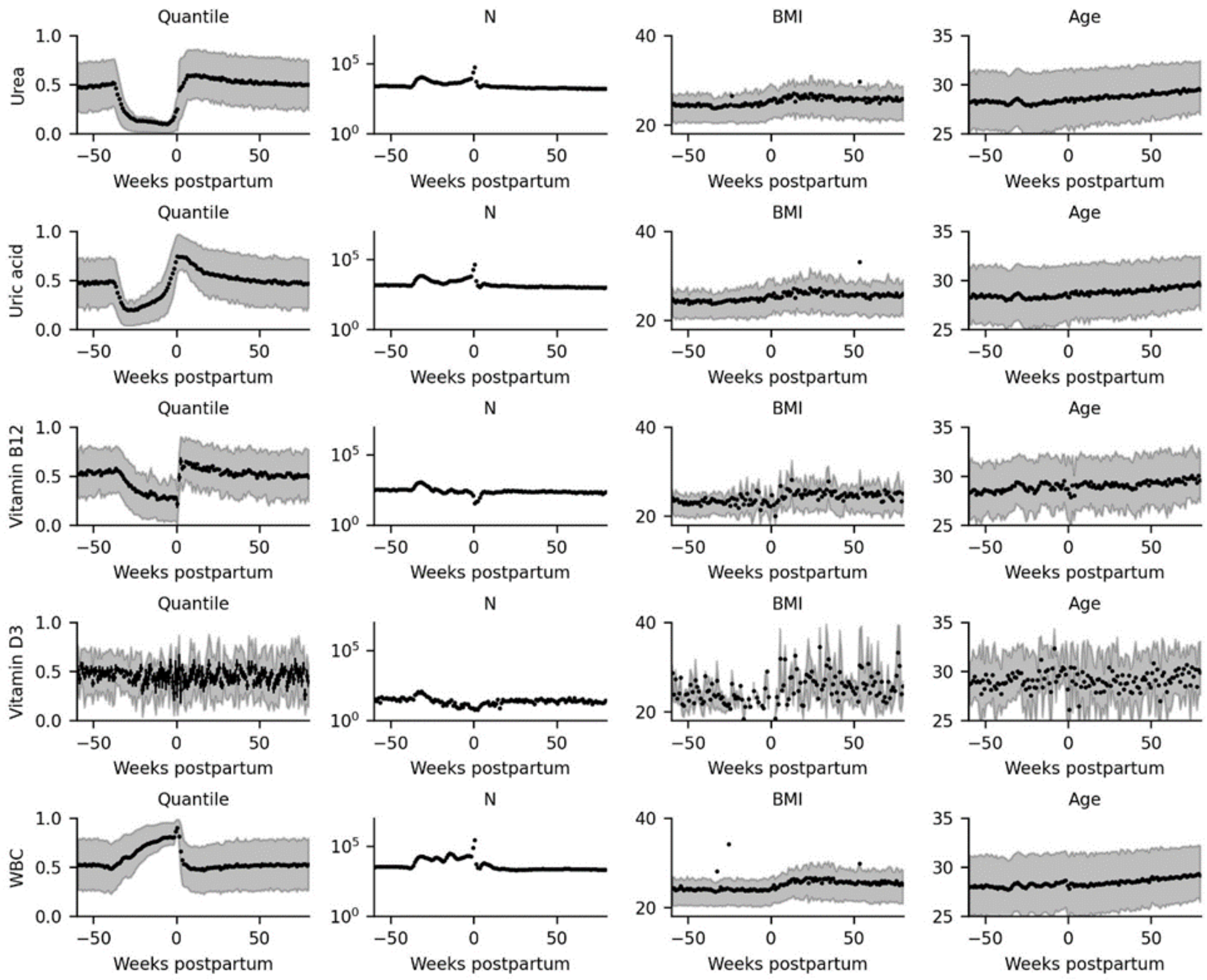


**110 lab test summary statistics in pregnancy and postpartum.** Each row represents a different lab test. Columns are (left to right): (Black) Mean percentile score at each 1-week interval, error bars (standard error of the mean) are usually smaller than the marker, (gray) IQR of percentile score. Number of measurements at each 1-week interval. (Black) Mean BMI of the population at the corresponding weekly interval, (gray) IQR of BMI. (Black) Mean age of the population at the corresponding weekly interval, (gray) IQR of age.

**Fig. S2.**

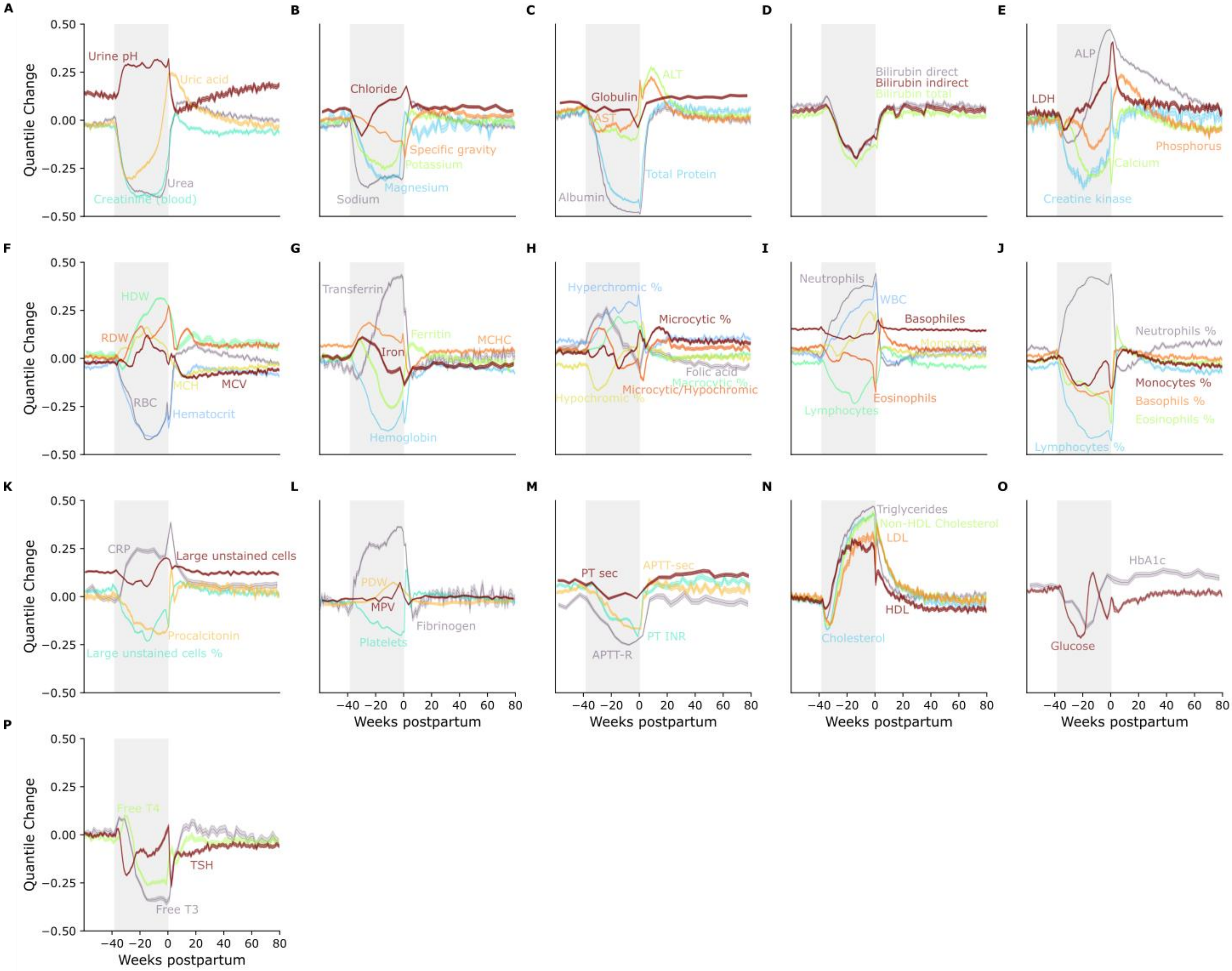


**Test dynamics, pregnancy is in gray.** Standard error of the mean is the thin lighter area around each line. For each system, the quantile score was used. All tests in the early preconception period hovered around the median. To show the relative change, 0.5 was subtracted. Some graphs were smoothed for visualization purposes (Methods).

**Fig. S3.**

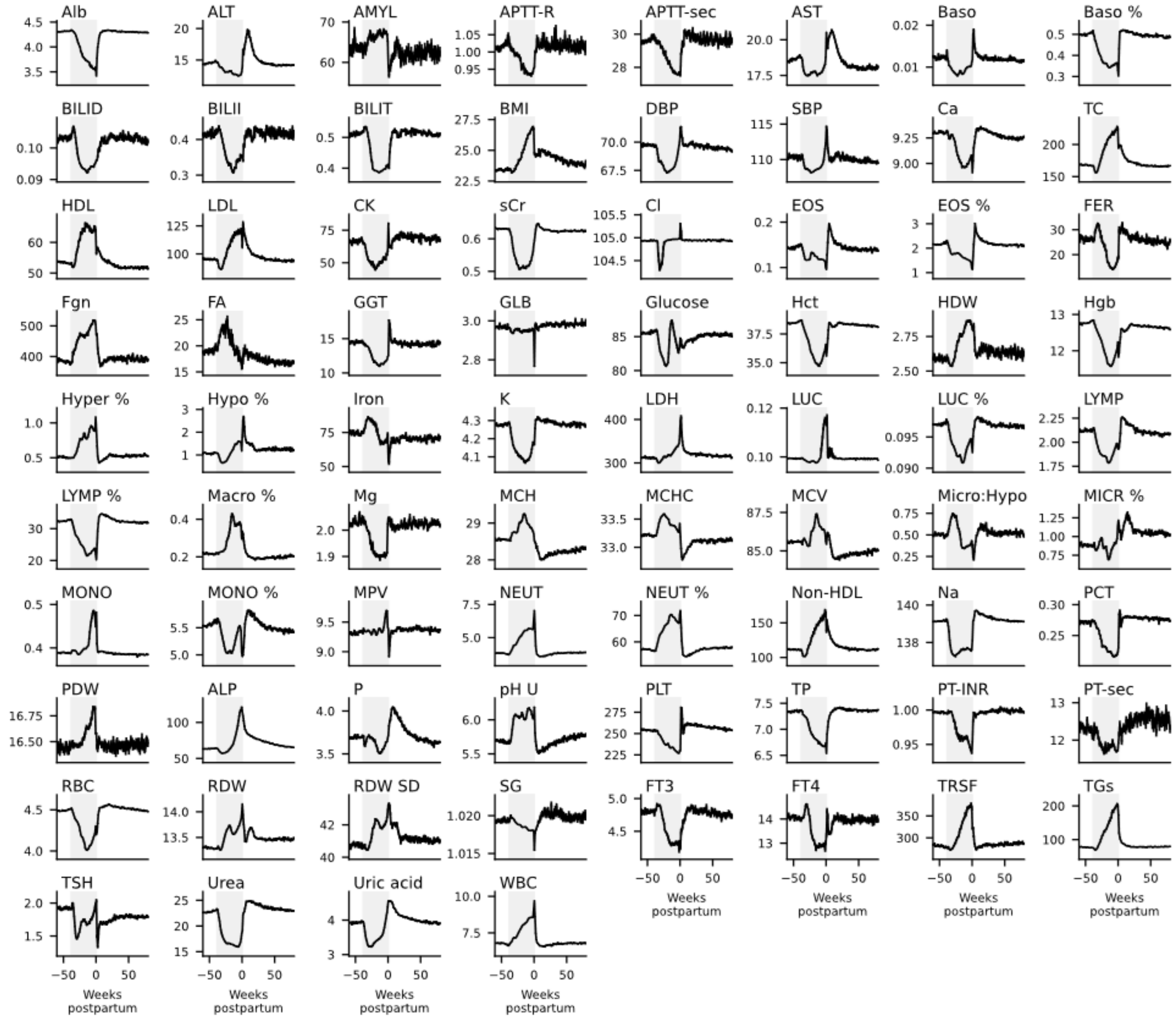


Back-transformed values of 76 tests over 140 weeks, no smoothing.

**Fig. S4.**

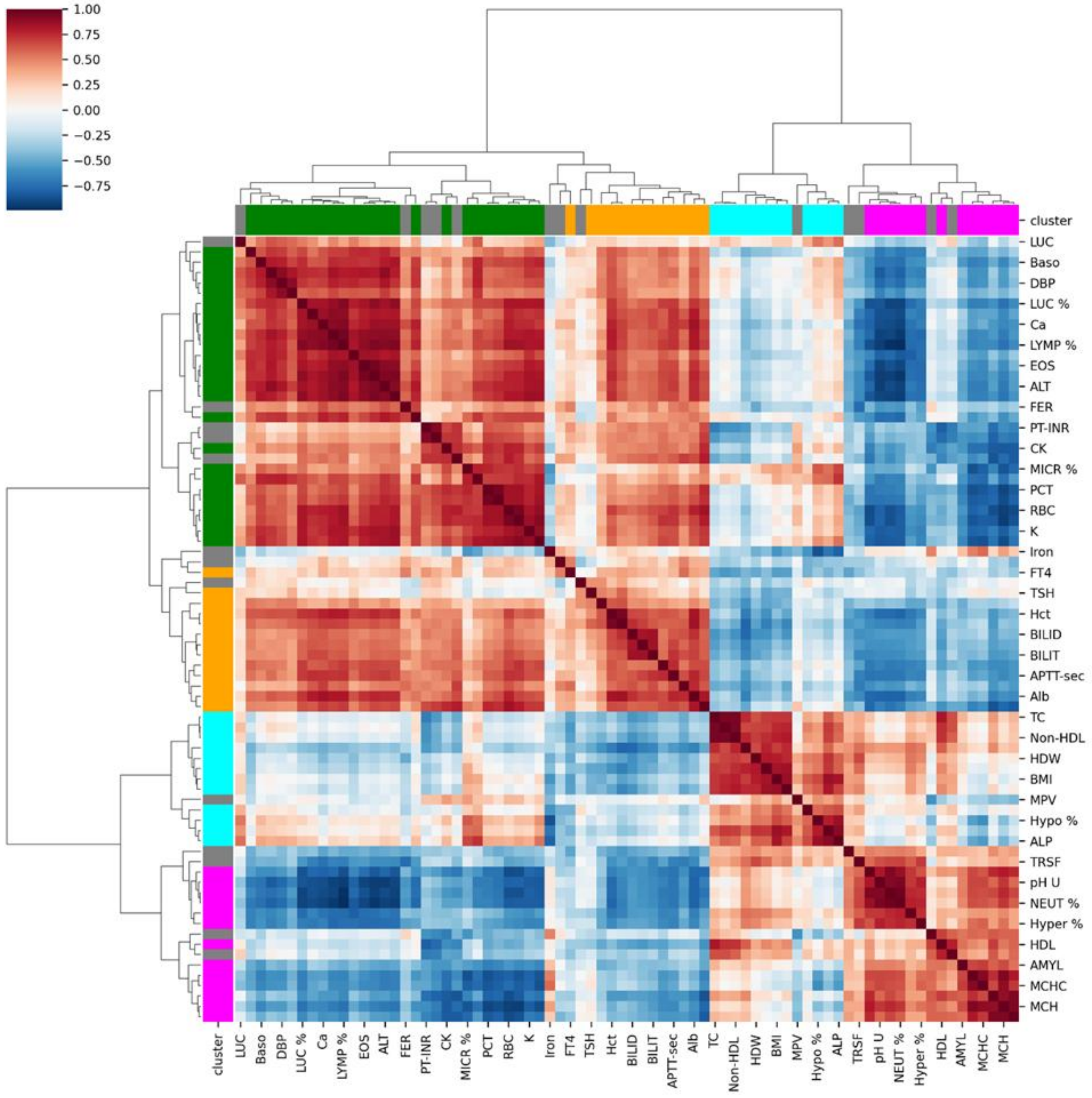


**Heatmap and dendrogram of clusters of tests.** Red/blue colormap is the distance between tests. Colors near the dendrograms denote clusters, gray denotes outlier tests which are more than one standard deviation away from the mean with respect to the above metric. Outliers are detected per cluster.

**Fig. S5.**

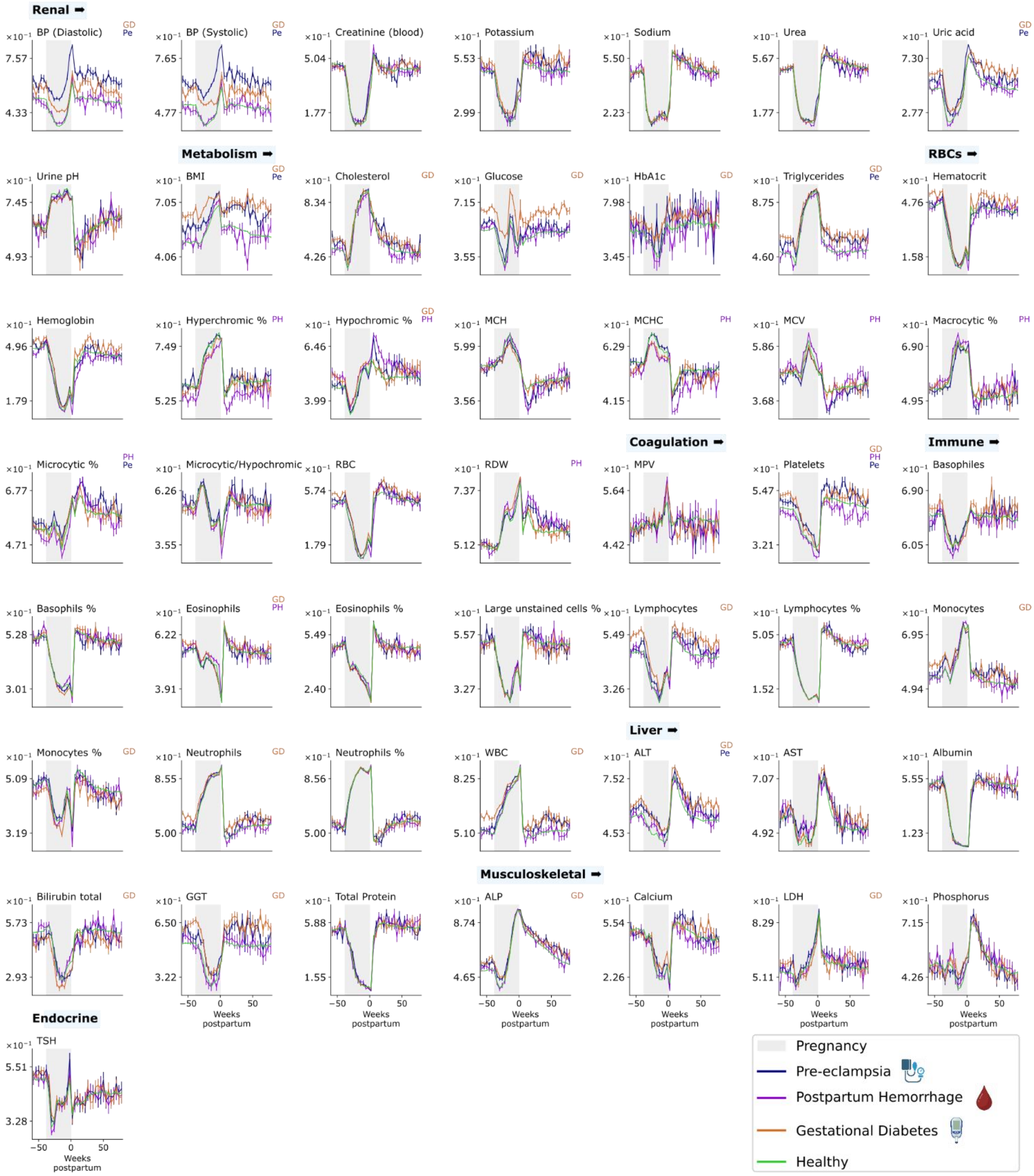


**Dynamics of lab tests in complications of pregnancy.** Y-axis is the quantile score.

**Table S1.**

| Name | File name | Abbreviation | LabNorm name | Group |
|---|---|---|---|---|
| 17α-OHP | 17_HYDROXY_PROGESTERONE | 17α-OHP | | Endocrine |
| ACTH | ACTH_ADRENOCORTICOTROPIC_HORMONE | ACTH | | Endocrine |
| Albumin | ALBUMIN | Alb | Albumin | Liver |
| Aldosterone | ALDOSTERONE_BLOOD | ALD | | Endocrine |
| ALT | ALT_Alanine_aminotransferase_GPT | ALT | ALT | Liver |
| Amylase | AMYLASE_BLOOD | AMYL | Amylase | Metabolism |
| AND | ANDROSTENEDIONE | AND | | Endocrine |
| ACE | ANGIOTENSIN_I_CONVERTING_ENZYME | ACE | | Endocrine |
| APTT-R | APTT_R | APTT-R | aPTT, ratio | Coagulation |
| APTT-sec | APTT_sec | APTT-sec | aPTT, sec | Coagulation |
| AST | AST_Aspartate_aminotransferase_GOT | AST | AST | Liver |
| Basophils % | BASO_perc | Baso % | Basophils, % | Immune |
| Basophiles | BASOPHILES_abs | Baso | Basophils, Abs | Immune |
| Bilirubin direct | BILIRUBIN_DIRECT | BILID | Direct Bilirubin | Liver |
| Bilirubin indirect | BILIRUBIN_INDIRECT | BILII | Indirect Bilirubin | Liver |
| Bilirubin total | BILIRUBIN_TOTAL | BILIT | Total Bilirubin | Liver |
| BP (Diastolic) | BPd | DBP | Blood Pressure, Diastolic | Renal |

| | | | | |
|---|---|---|---|---|
| BP (Systolic) | BPs | SBP | Blood Pressure, Systolic | Renal |
| BMI | BMI | BMI | BMI | Metabolism |
| CRP | C_REACTIVE_PROTEIN_CRP | CRP | CRP | Immune |
| Calcium | CALCIUM_BLOOD | Ca | Calcium | Musculoske letal |
| Cholesterol | CHOLESTEROL | CL | Total Cholesterol | Metabolism |
| HDL | CHOLESTEROL_HDL | HDL | HDL Cholesterol | Metabolism |
| HDL ratio | CHOLESTEROL_HDL_RATIO | HDL ratio | Total:HDL | Metabolism |
| LDL | CHOLESTEROL_LDL | LDL | LDL Cholesterol | Metabolism |
| Creatine kinase | CK_CREAT | CK | CPK | Musculoske letal |
| Chloride | Cl | Cl | Chloride | Renal |
| Cortisol | CORTISOL_BLOOD | Cort | | Endocrine |
| Cortisol urine 24h | CORTISOL_U_FREE_24h | Cort 24h | | Endocrine |
| Cortisol urine free samp | CORTISOL_U_FREE_SAMP | Cort u samp | | Endocrine |
| Creatinine (blood) | CREATININE_BLOOD | sCr | Creatinine | Renal |
| Creatinine (urine) | CREATININE_URINE_SAMPLE | uCr | Urine Creatinine | Renal |
| DHEA-S | DHEA_SULPHATE | DHEA-S | | Endocrine |
| Eosinophils | EOS | EOS | Eosinophils, Abs | Immune |
| Eosinophils % | EOS_perc | EOS % | Eosinophils, % | Immune |
| Estradiol | ESTRADIOL_E_2 | E2 | Estradiol | Endocrine |
| Ferritin | FERRITIN | FER | Ferritin | RBCs |
| Fibrinogen | FIBRINOGEN | Fgn | Fibrinogen | Coagulation |
| Folic acid | FOLIC_ACID | FA | Folic Acid | RBCs |

| | | | | |
|---|---|---|---|---|
| FSH | FSH_FOLLICLE_STIMULATING_HORMONE | FSH | | Endocrine |
| GGT | GAMMA_GLUTAMYL_TRANSPEPTIDASE | GGT | GGT | Liver |
| Globulin | GLOBULIN | GLB | Total Globulin | Liver |
| Glucose* | GLUCOSE_BLOOD | Glucose | Glucose | Metabolism |
| Growth Hormone | GROWTH_HORMONE_GH | GH | | Endocrine |
| Hematocrit | HCT | Hct | Hematocrit | RBCs |
| Hct/Hgb | HCT_HGB_RATIO | Hct/Hgb | Hematocrit:Hemoglobin | RBCs |
| HDW | HDW | HDW | HDW | RBCs |
| HbA1c | HEMOGLOBIN_A1C_CALCULATED | HbA1c | Hemoglobin A1c | Metabolism |
| Hemoglobin | HGB | Hgb | Hemoglobin | RBCs |
| Hyperchromic % | HYPERperc | Hyper % | Hyperchromic | RBCs |
| Hypochromic % | HYPO_perc | Hypo % | Hypochromic | RBCs |
| Insulin | INSULIN | Insulin | | Metabolism |
| IGF-1 | INSULIN_LIKE_GROWTH_FACTOR_1 | IGF-1 | | Endocrine |
| Iron | IRON | Iron | Iron | RBCs |
| Potassium | K | K | Potassium | Renal |
| LDH | LACTIC_DEHYDROGENASE_LDH__BLOOD | LDH | LDH | Musculoskeletal |
| LH | LH_LUTEINIZING_HORMONE | LH | LH | Endocrine |
| Large unstained cells | LUC | LUC | Large unstained cells, Abs | Immune |

| | | | | |
|---|---|---|---|---|
| Large unstained cells % | LUCperc | LUC % | Large unstained cells, % | Immune |
| Lymphocytes | LYMP | LYMP | Lymphocytes, Abs | Immune |
| Lymphocytes % | LYMperc | LYMP % | Lymphocytes, % | Immune |
| Microalbumin/Creatinine | M | ACR | Microalbumin:Creatinine | Renal |
| Macrocytic % | MACROperc | Macro % | Macrocytic | RBCs |
| Magnesium | MAGNESIUM_BLOOD | Mg | Magnesium | Renal |
| MCH | MCH | MCH | MCH | RBCs |
| MCHC | MCHC | MCHC | MCHC | RBCs |
| MCV | MCV | MCV | MCV | RBCs |
| Microcytic % | MICR_perc | MICR % | Microcytic | RBCs |
| Microalbumin urine 24h | MICROALBU_U_SAMP | UMA | Urine Microalbumin | Renal |
| Microcytic/Hypochromic | MICROperc_HYPOperc | Micro:Hypo | Microcytic:Hypochromic | RBCs |
| Monocytes | MONO | MONO | Monocytes, Abs | Immune |
| Monocytes % | MONperc | MONO % | Monocytes, % | Immune |
| MPV | MPV | MPV | MPV | Coagulation |
| MPXI | MPXI | MPXI | MPXI | Immune |
| Sodium | Na | Na | Sodium | Renal |
| Neutrophils | NEUT | NEUT | Neutrophils, Abs | Immune |
| Neutrophils % | NEUTperc | NEUT % | Neutrophils, % | Immune |
| Non-HDL Cholesterol | NON_HDL_CHOLESTEROL | Non-HDL | Non-HDL | Metabolism |
| PTH | PARATHYROID_HORMONE_PTH | PTH | | Musculoskeletal |
| Procalcitonin | PCT | PCT | PCT | Immune |

| | | | | |
|---|---|---|---|---|
| PDW | PDW | PDW | PDW | Coagulation |
| Urine pH | PH_u | pH U | Urine pH | Renal |
| Alkaline Phosphatase | PHOSPHATASE_ALKALINE | ALP | Alk. Phosphatase | Musculoskeletal |
| Phosphorus | PHOSPHORUS_BLOOD | P | Phosphorus | Musculoskeletal |
| Platelets | PLT | PLT | Platelets | Coagulation |
| Progesterone | PROGESTERONE | P4 | | Endocrine |
| Prolactin | PROLACTIN | PRL | | Endocrine |
| Total Protein | PROTEIN_TOTAL_BLOOD | TP | Total Protein | Liver |
| PT INR | PT_INR | PT-INR | PT, INR | Coagulation |
| PT % | PT_perc | PT % | PT, % | Coagulation |
| PT sec | PT_SEC | PT-sec | PT, sec | Coagulation |
| RBC | RBC | RBC | RBC | RBCs |
| RBC u strip | RBC_calculated_urine_strip | RBC u strip | | Renal |
| RDW | RDW | RDW | RDW | RBCs |
| RDW SD | RDW_SD | RDW SD | RDW-SD | RBCs |
| Sedimentation rate | SEDIMENTATION_RATE | Sed Rate | ESR | Immune |
| Specific gravity | SPECIFIC_GRAVITY | SG | Urine Specific Gravity | Renal |
| Free T3 | T3_FREE | FT3 | T3, Free | Endocrine |
| Total T3 | T3_TOTAL | TT3 | | Endocrine |
| Free T4 | T4_FREE | FT4 | T4, Free | Endocrine |
| Testosterone | TESTOSTERONE_TOTAL | T | | Endocrine |
| Thyroglobulin | THYROGLOBULIN | Tg | | Endocrine |
| Transferrin | TRANSFERRIN | TRF | Transferrin | RBCs |
| Triglycerides | TRIGLYCERIDES | TGs | Triglycerides | Metabolism |

| | | | | |
|---|---|---|---|---|
| TSH | TSH_THYROID_STIMULATING_HORMONE | TSH | TSH | Endocrine |
| Urea | UREA_BLOOD | Urea | Urea | Renal |
| Uric acid | URIC_ACID_BLOOD | Uric acid | Uric Acid | Renal |
| Vitamin B12 | VITAMIN_B12 | VitB12 | Vitamin B12 | RBCs |
| Vitamin D3 | VITAMIN_D3_25_0H_RIA | VitD3 | Vitamin D (25-OH) | Musculoskeletal |
| WBC | WBC | WBC | WBC | Immune |

* Glucose test – fasting glucose test. Instructions for the test include an 8 hour fast before drawing blood. No limitation on hydration.

**Laboratory tests in the dataset**. In **red** are the tests which had a high level of noise/low number of measurements. These tests were excluded out from analyses, except for HBA1c was re-introduced in the analyses for the **"Complications in Pregnancy"** section due to its importance in gestational diabetes. CRP was included in the section **"Health style behaviors are reflected in preconception dynamics"** for its importance and low noise during the preconception period. In **blue** are additional tests which were excluded from the analyses in the "Complications in Pregnancy" section for the same reasons due to the smaller number of participants.

**Table S2.**

| | |
|---|---|
| $i, i \in [-60,80]$ | The relative week of the measurement, where 0 is delivery, in weekly resolution. |
| $\mu\_\{x\_i\}, \sigma\_\{x\_i\}, n\_\{x\_i\}$ | The mean/standard deviation/number of all measurements of test $x$ at week $i$. |
| $\lfloor \alpha_{\{x_i\}} \rfloor$ | The floor of the median age of the cohort for test $x$ at week $i$ |
| ${x_{(q)}}_i, q \in (0,1)$ | The $q^{th}$ quantile of lab test $x$ at week $i$, compared to healthy, non-pregnant same aged, reference population (**Methods**). |
| $\mu^q_{x_i}, \sigma^q_{x_i}$ | The mean/standard deviation of quantile score of test $x$ at week $i$. |
| $F_x^{-1}: (0,1) \times N \rightarrow Range(x)$ | The back-transform of test $x$ from the quantile score back to the test values (**Methods**). First argument is the quantile score, second is the age for the back-transformation. |
| $\widetilde{\mu_{x_\iota}} = F_x^{-1}\left(\mu^q_{x_i}, \lfloor \alpha_{x_i} \rfloor\right)$ | Shorthand for the back-transform of the mean quantile score of test $x$ at week $i$. |

Notations for noise threshold calculation.

**Table S3.**

| | Number of measurements lower threshold $\left(n_{x_i}\right)$ | | Noise upper threshold $\left(\zeta_{x_i} = \frac{\sigma_{i}{}^{err}_{F_x^{-1}}}{\sigma_{F_x^{-1}}}\right)$ | |
|---|---|---|---|---|
| | $\leq \min\left\{n_{x_i}\right\}_i$ (Any weekly interval) | $\leq \overline{n_{x_i}}$ (Mean across all weekly intervals) | $\leq \max\left\{\zeta_{x_i}\right\}_i$ (Any weekly interval) | $\leq \overline{\zeta_{x_i}}$ (Mean across all weekly intervals) |
| General Dataset | 50 | 80 | 0.7 | 0.5 |
| Complications dataset | 10 | 80 | 1.3 | 1.0 |

**Thresholds for filtering the dataset.** Tests with fewer measurements than stated were excluded, tests with greater noise level than stated were excluded too (except HBA1c as noted). See supplementary ("Noise Filtering") the explanation for $\sigma_{i}{}^{err}_{F_x^{-1}}$ and $\sigma_{F_x^{-1}}$. Thresholds were set by eye for the general dataset to retain results with reasonably small fluctuations between weeks and small enough standard deviation of the mean. Thresholds for the complications dataset are slightly more lenient due to the lower number of participants and higher variance.
